# Coarse-graining metabolic networks via feature learning reveals cross-species growth predictors

**DOI:** 10.64898/2026.05.13.725055

**Authors:** Aoyu Zhu, Po-Yi Ho

## Abstract

Bacterial growth and the underlying metabolic networks are highly dissimilar across species, posing a fundamental challenge for various bioengineering tasks. For a given species across nutrient environments, growth is regulated via proteome allocation, which gives rise to linear relationships between growth and the fractions of coarse-grained proteome sectors. However, it remains unclear whether there are coarse-grained predictors that correlate with growth across species monocultures in a given environment. Here, simulating genome-scale metabolic models, we discover two conserved growth predictors: the number of nutrients utilized, and a set of metabolic sectors that we identify by coarse-graining metabolic networks via feature learning. We find evidence supporting both predictors in recent experiments; notably, the proteome fractions of the sectors correlate with growth across species in a gene expression dataset. We further analyze coarse-grained networks to show that interactions between sectors improve growth predictions. Our work provides a framework to investigate how coarse-grained growth predictors emerge across diverse metabolic networks.

## INTRODUCTION

Bacterial growth depends on metabolic networks whose structures are highly dissimilar across species^1–4^. This dissimilarity poses a fundamental challenge for bioengineering where there is an increasing need to manipulate diverse species^5,6^. In addition to species-specific network structures, growth is also determined by how fluxes over metabolic reactions are regulated in response to nutrient environments^7–10^. For model organisms in minimal environments, growth regulation can be understood using simple quantitative rules with broad implications for synthetic biology^11^. However, the same is not true for more diverse contexts. While there exist computational tools to predict growth given metabolic networks and nutrient environments^12,13^, their utility has been limited by a lack of simple rules to reason about growth differences. Here, we address this gap by finding coarse-grained features of metabolic networks that predict growth across species (**Figure 1**).

**Figure 1.**
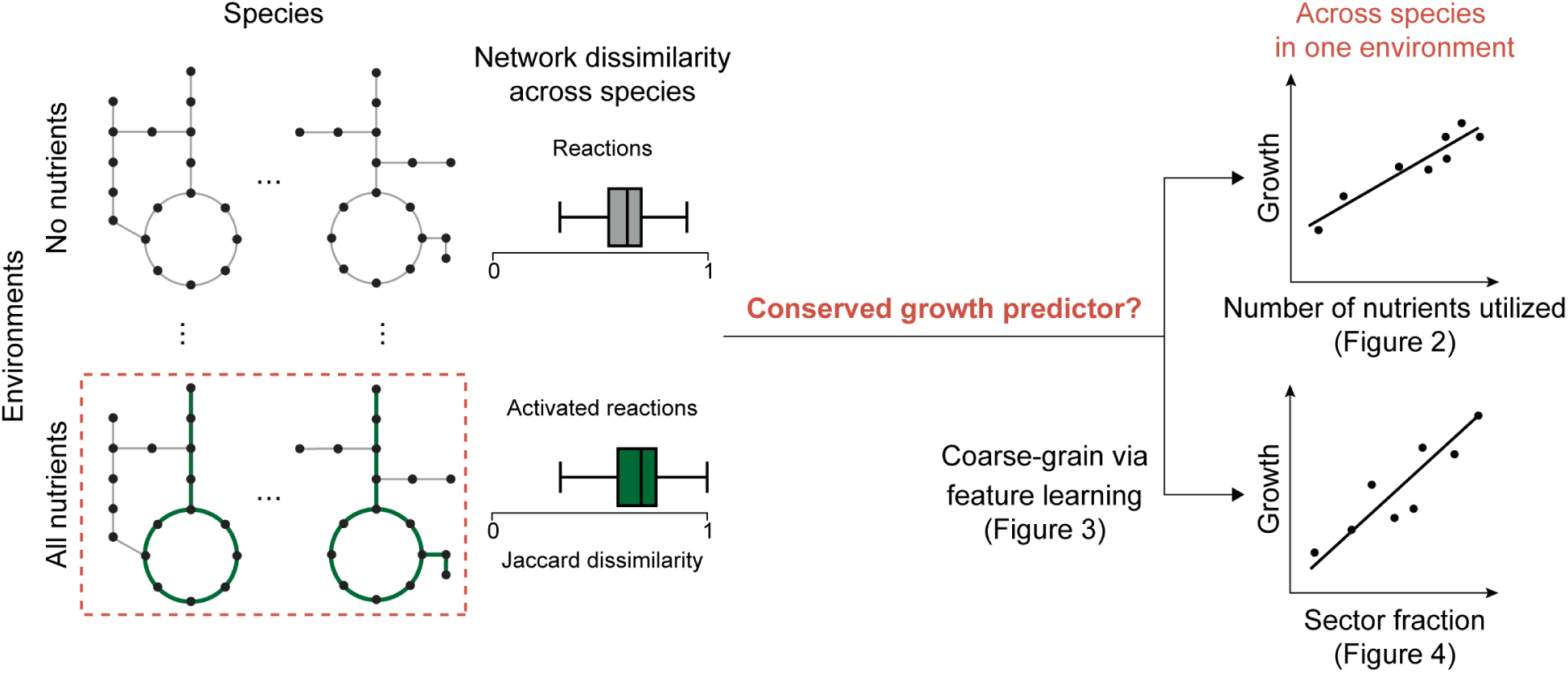
Are there simple growth predictors that are conserved across species despite dissimilar metabolic networks?

For *E. coli* grown in defined media with only one carbon source, growth rate is related to the supplied concentration of the carbon source by the Monod equation^7^. In other words, the uptake flux is a simple growth predictor. On two carbon sources, the regulatory choice between co-utilization and diauxie again depends on the uptake flux^14^. Importantly, this regulatory choice is predicted by proteome allocation^15^, the constraint that expressing a metabolic pathway takes away some of the limited cellular resources from other pathways. Proteome allocation gives rise to empirical trends known as bacterial growth laws, linear relationships between growth rate and the fractions of proteome sectors for a given species across environments^11^. At the coarsest level, growth rate scales linearly with the ribosomal fraction of the proteome for *E. coli* grown on various carbon sources^8,16^. Other sectors, such as groups of genes involved in catabolism, also contribute to and scale with growth^17^. These growth laws are active consequences of regulatory strategies that sense reaction fluxes to balance the proteome cost of a sector with its growth benefit^18–24^. As a result, proteome sectors, when appropriately defined, are also simple growth predictors.

But are there growth predictors that are conserved across species? On top of the fact that metabolic networks are dissimilar across species, regulatory strategies and the resulting growth outputs can become complex already for a small number of nutrients^25,26^, even though natural contexts such as human guts can contain many more^27^. As a result, it remains unclear whether there are proteome sectors that predict growth across species, and if so, how to identify them in general. Encouragingly, cross-environment ribosomal growth laws have recently been observed in several other species^28^, suggesting a wider search for conserved growth predictors.

To do so, we sought to extract coarse-grained features from metabolic networks – i.e., quantities that are derived from network structures and that involve groups of metabolic reactions. However, existing methods to coarse-grain metabolic networks are not suitable for our purposes. For example, reactions can be grouped according to annotations such as metabolic subsystem classifications^29^ or via clustering expression data^17,30,31^, but these methods are hard to generalize across species or environments for which data are lacking. Alternatively, flux networks can be decomposed without relying on additional data via enumerating network paths, but most existing methods are combinatorially explosive^32^, lack resolution^33^, or require implicit undefined components^34^. To overcome these challenges, we first simulated metabolic networks under a simple proteome constraint model to predict reaction fluxes. This step could be efficiently implemented via linear programming, which allowed us to sample regulatory outputs across species and environments. Then, we used feature learning to embed the diverse flux networks without enumerating exact paths, thereby identifying reactions that are under similar regulation across contexts.

We applied this approach to an ensemble of diverse metabolic networks and found two conserved growth predictors: the number of nutrients utilized (**Figure 2**) and a set of metabolic sectors (**Figure 3-4**). Both predictors exhibited significant cross-species correlations with growth in several recently published experiments. These results provide multiple, independent lines of evidence and a theoretical framework for further investigations into conserved growth predictors.

**Figure 2.**
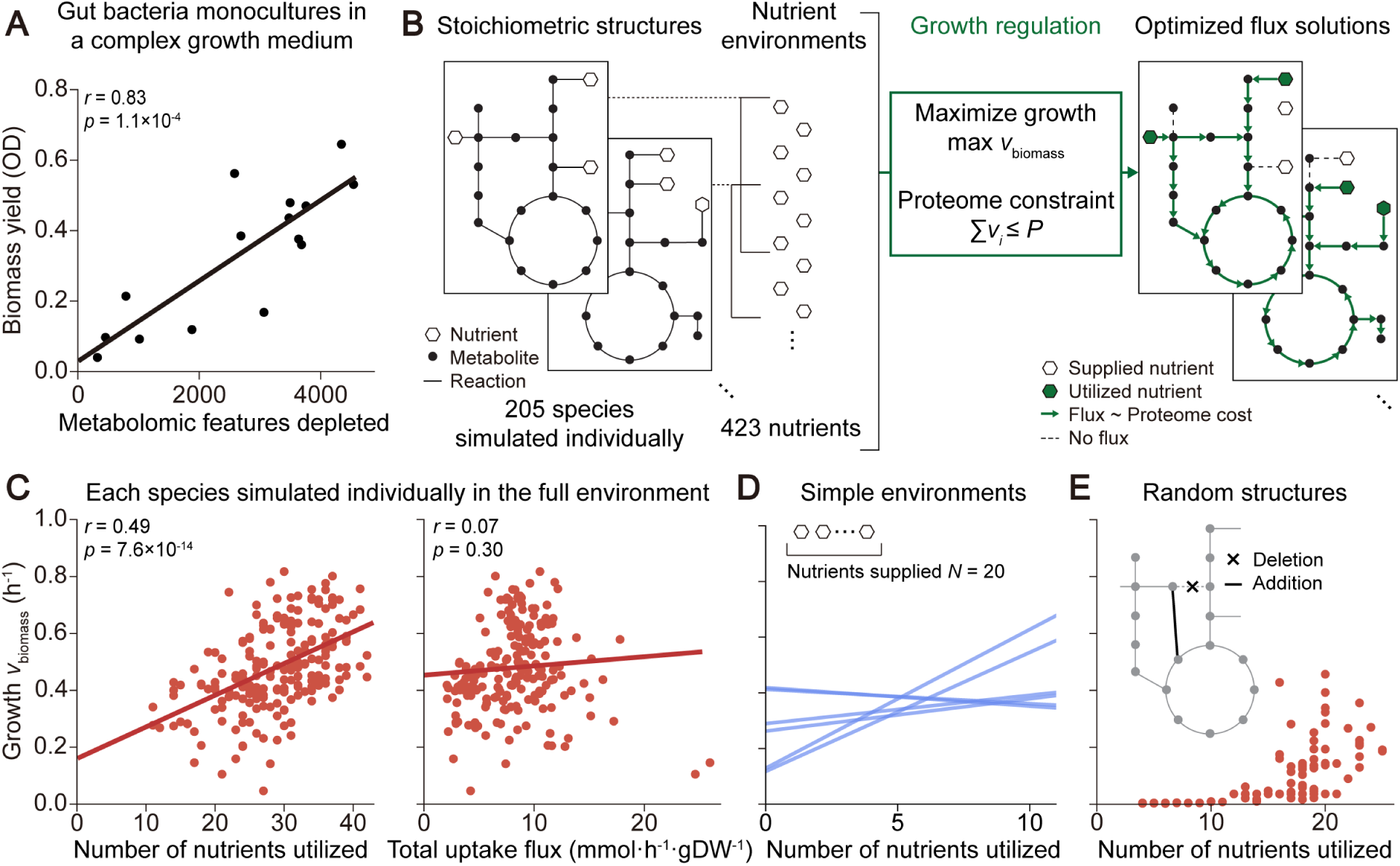
The number of nutrients utilized correlates with growth across diverse metabolic networks. A) Biomass yield was well correlated with the number of metabolomic features depleted for 15 gut bacterial species grown in monoculture in a complex medium (Pearson’s *r* = 0.83, *p* = 1e−4; reproduced from Ref.^35^). The line shows the best linear fit. Metabolomic features represent nutrients in the environment; one nutrient can generate multiple metabolomic features. B) Simulating metabolic networks under proteome-constrained growth maximization (**Methods**). Optimized flux solutions were obtained via flux balance analysis, taking as inputs the stoichiometric structure of a species and the set of nutrients that make up the environment. Stoichiometric structures of 205 phylogenetically diverse bacterial species were obtained from the AGORA2 database of genome-scale metabolic models^37^. Species can uptake different sets of nutrients as specified by the available exchange reactions, and the total set of all non-drug exchange reactions across species made up 423 nutrients. A proteome constraint was implemented by constraining the sum of fluxes to be less than a constant value fixed across species. Each species was simulated individually, without interactions from other species. After growth regulation, only some nutrients are utilized and only some reactions are activated. C) Growth was correlated with the number of nutrients utilized, across species simulated individually in the full environment with all 423 potential nutrients supplied (*r* = 0.49, *p* = 8e−14). Growth was quantified via flux through the biomass reaction, scaled such that the unit can be approximately interpreted as per hour (**Methods**). By contrast, growth was not correlated with the total uptake flux of nutrients (*r* = 0.07, *p* = 0.30). D) The correlation between growth and the number of nutrients utilized was highly variable across simple environments each containing 20 randomly selected nutrients (**Methods**). Each line shows the best fit linear regression across species in one environment. E) Growth was not well correlated with the number of nutrients utilized for an ensemble of biologically implausible networks, generated from the *E. coli* model by randomly deleting and adding reactions (**Methods**).

**Figure 3.**
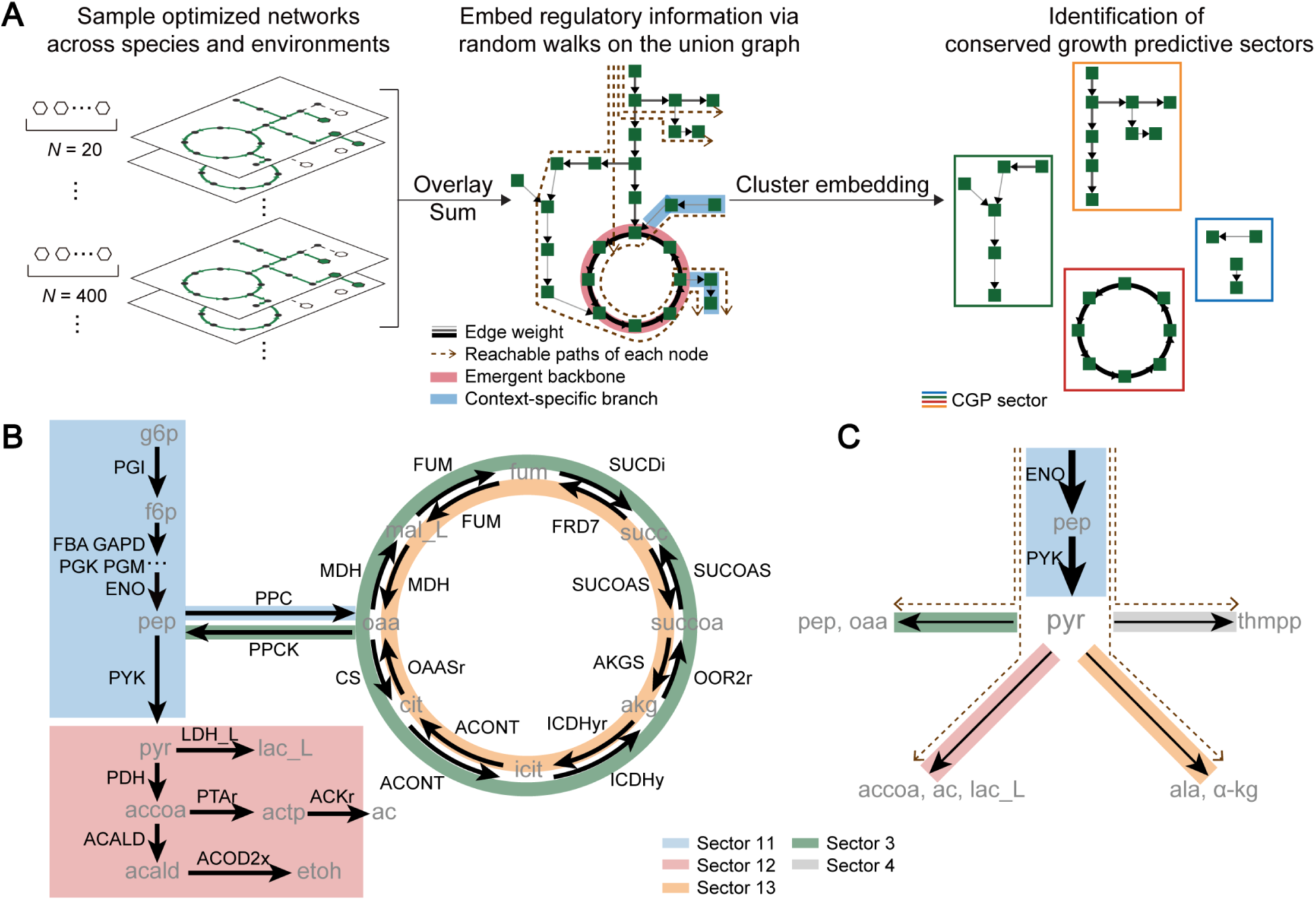
Coarse-graining metabolic networks. A) Identifying coarse-grained network features that emerge from shared growth regulation (**Methods**). Optimized networks in the form of reaction-based graphs (**Figure S3**), sampled across species and environments, were overlaid and summed to obtain a union graph. Edge weights in the union graph captured usage frequencies. High-weight edges revealed emergent backbones shared across contexts, and low-weight edges corresponded to context-specific branches. Reactions were embedded into a latent space using node2vec to sample reachable paths for each node via random walks on the union graph^50^. The embeddings were clustered using *k*-means to obtain groups of reactions under similar regulation across diverse contexts. This procedure resulted in 20 reaction groups that we refer to as conserved growth predictive (CGP) sectors. B) CGP sectors captured functional information in central carbon metabolism. Shown here are glycolysis, the TCA cycle, and fermentation. Colors indicate reaction-sector assignments. C) CGP sectors captured flux-dependent network features. Shown are reachable paths passing through pyruvate, which predominantly originated from sector 11 and branched into several downstream sectors.

**Figure 4.**
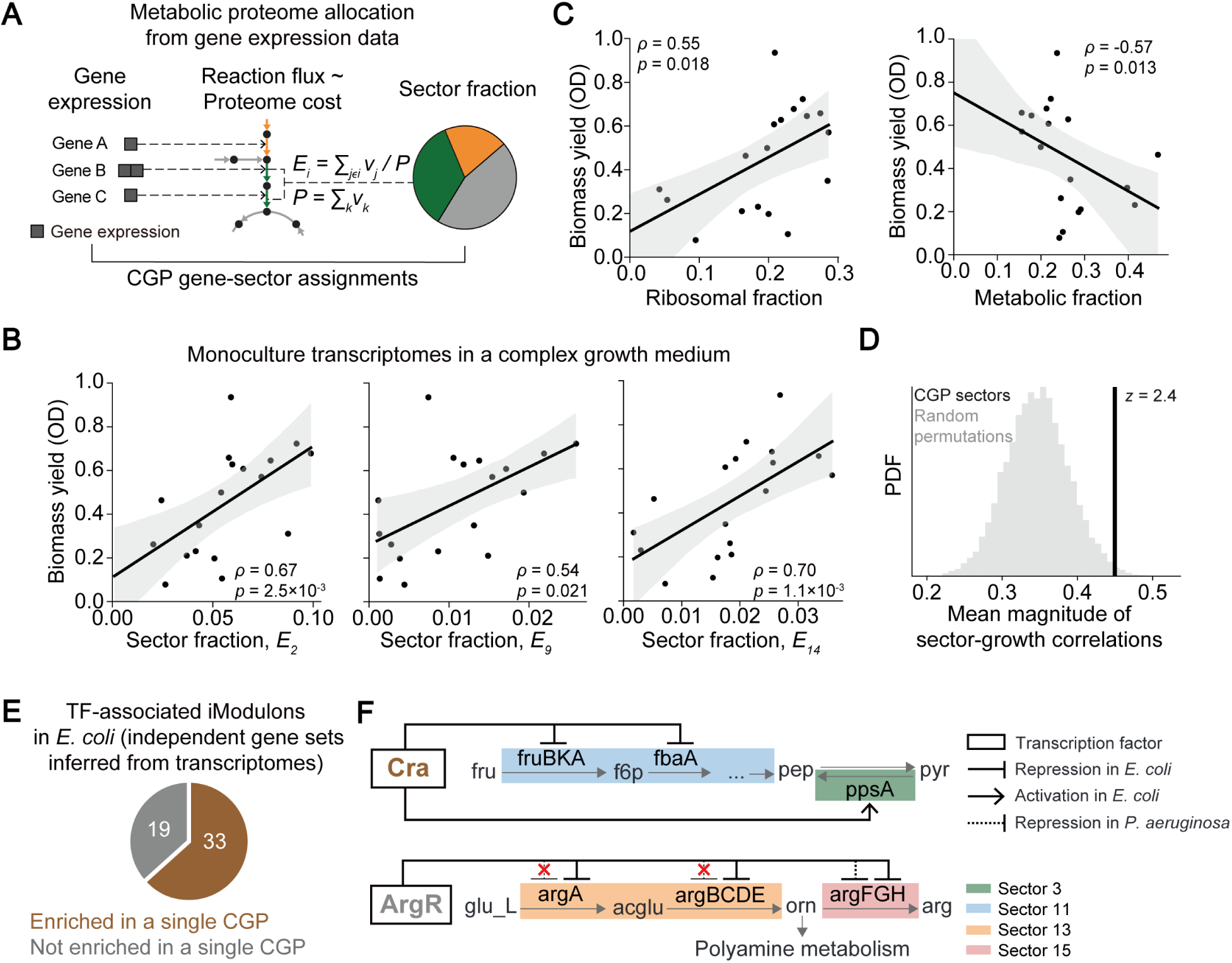
Sector fractions correlate with growth across species monocultures in a transcriptome dataset. A) Quantifying metabolic proteome allocation from gene expression data. The expression level of a gene was assumed to be proportional to the flux and proteome cost of the reaction that it encodes. If a gene encodes multiple reactions, its expression level was evenly distributed across the encoded reactions. Sector fractions were calculated by dividing by the total proteome cost over all reactions (**Methods**). B) (B-D) Transcriptome and biomass yield data from Ref.^52^ were analyzed. The data were obtained for 18 gut bacterial species grown in monoculture in a complex medium. Shown are correlations between growth and sector fractions for 3 example sectors (Spearman’s *ρ* > 0.54 and *p* < 0.02 for all 3 sectors; see **Figure S6** for all sectors). Solid lines and shaded areas indicate linear fits and 95% confidence intervals. C) The ribosomal and total metabolic fractions were also correlated with growth (*ρ* = 0.55 and −0.57, *p* = 0.02 and 0.01). D) Growth correlations of the CGP sectors were not statistical artifacts. Overall performance was measured by the average over sectors of the magnitude of growth correlations with sector fractions. Shown is the distribution of this quantity for random permutations of reaction-sector assignments that preserve the number of reactions in each sector, compared to the value for the CGP assignment (*z*-score = 2.4; **Methods**). E) Genes that are putatively co-regulated by the same transcription factor – i.e., those assigned to the same transcription factor-associated iModulon via independent component analysis on *E. coli* transcriptome data^31^ – tended to also be grouped together in the same CGP sector (**Methods**). F) CGP sectors can be implemented by transcription factors. Top: Cra represses fruBKA and fbaA (assigned to sector 11) but activates ppsA (sector 3). Bottom: ArgR represses argA-E in *E. coli* but not in *P. aeruginosa* (sector 13), and represses argFGH in both species (sector 15).

## RESULTS

### Proteome-constrained optimization predicts cross-species trends in growth and nutrient utilization

A recent experiment found that biomass yield was strongly correlated to the number of nutrients utilized across 15 gut bacteria grown individually in a complex growth medium^35^ (*r* = 0.83, *p* = 1e−4; **Figure 2A**). Inspired by this finding, which was unexplained, we searched in silico for cross-species trends in growth and nutrient utilization. We reasoned that cross-species trends can emerge if the species use approximately the same growth regulatory strategy, which we hypothesized to be growth maximization under a proteome constraint^24^. To test this hypothesis, we used flux balance analysis (FBA)^12^, which takes as input a stoichiometric matrix that describes the structure of the metabolic network of a species and a set of nutrients supplied that describes the environment. FBA then solves a constrained optimization problem to output a flux distribution that simultaneously maintains the network at steady state and maximizes growth rate, as modeled by the flux through a biomass reaction. We implemented a simplified proteome constraint by assuming that the proteome cost of a reaction is proportional to its flux and that the total cost across all reactions must be less than a constant value fixed across species. We further assumed that nutrients were supplied at large enough amounts so that the proteome constraint is the growth limiting factor (**Methods**, **Figure 2B**), as is the case in typical laboratory experiments^36^. Since stoichiometric matrices can be constructed just from genomes, this framework allowed us to investigate cross-species trends despite a lack of experimental data.

We applied this framework to analyze a collection of high-quality metabolic models of 205 phylogenetically diverse species^37^. These models contained dissimilar sets of reactions, even when considering only reactions with flux, when considering only reactions in central metabolic subsystems, and when accounting for slight variants in reactants (**Figure 1, S1**). In the full environment where all possible nutrients were supplied, the predicted growth was significantly correlated to the number of nutrients utilized *M* across species (*r* = 0.49, *p* = 8e−14; **Figure 2C**), in qualitative agreement with the experimental observation.

We note the caveat that biomass yield and growth rate are different quantities. However, they both depend on the underlying metabolic network, and as shown in several recent work, can be correlated both experimentally and theoretically in certain contexts^38,39^. Moreover, the biomass pseudo-reaction in FBA is in any case a theoretical construct that is only indirectly linked to either rate or yield. As such, FBA is often used empirically to predict both yield and rate^12,40–43^. Also, we did not explicitly link models and experiments for each species individually. Nonetheless, stoichiometric structures are accurate in aggregate^37,44,45^, enabling analyses at the ensemble level. Thus, we interpret **Figure 2A** and **2C** to provide independent, qualitatively consistent evidence that *M* is a conserved growth predictor.

By contrast, in the same simulations, growth was not correlated to the total input flux nor its carbon fraction (*r* = 0.07 and 0.11, *p* = 0.30 and 0.13; **Figure 2C, S2**). This result implies that growth correlations with *M* were not a simple consequence of mass conservation, as expected given growth regulatory processes like overflow metabolism that result in secretions^46^. Active regulation was more important than static model properties in this prediction: while network size was correlated with growth (*r* = 0.38, *p* = 3e-8; **Figure S2**), this correlation was weaker than the correlation between growth and *M*, and network size was not correlated with *M* itself (*r* = -0.05, *p* = 0.44; **Figure S2**). These results were relatively robust to variations in the simplified proteome cost model, which we discuss in a later section.

The status of *M* as a conserved growth predictor depended on the nutrient environment and the network ensemble. Across simple environments where the number of nutrients supplied *N* was small (20 out of the 423 possible nutrients for the 205 species analyzed), growth correlations with *M* were highly variable and uncorrelated on average (**Figure 2D**). Growth was also uncorrelated with *M* for an ensemble of biologically implausible networks, which we generated by randomly adding and deleting reactions from an *E. coli* model while ensuring thermodynamic feasibility^47,48^ (**Methods**, **Figure 2E**). Thus, for realistic networks in complex environments, proteome-constrained optimization drove *M* to emerge as a conserved growth predictor.

### Coarse-graining flux networks links metabolic function to network features

Next, we searched for coarse-grained growth predictors involving more than just the outermost layer of metabolic networks, which required us to extract features from complex networks. In brief, we first re-casted flux networks into reaction-based graphs, since proteome costs are associated with reactions not metabolites. Simple path tracing on reaction-based graphs quantified how some nutrients contributed directly to biomass while others were multifunctional, demonstrating that reaction-based graphs highlight pathway-like features that might correlate with growth. Then, we sampled flux networks across species and environments, aggregated them into a union reaction-based graph, embedded the graph via feature learning, and clustered reactions into metabolic sectors (**Figure 3**). The resulting sectors captured canonical biochemical organizations, and changes in sector fractions on nutrient dropout correlated with whether the nutrients were multifunctional, demonstrating the interpretability of the learned sectors. We then tested the sectors on experimental data, which we discuss in the next section. Since the relationship between *M* and the sectors is unknown, we interpreted the analyses in the previous and next sections as largely independent.

In more detail, we wondered whether *M* increases growth by allowing nutrients to enter networks more downstream and closer to the biomass reaction, which should allow more input flux to contribute to growth at lower proteome costs from fewer intermediate reactions. To quantify this notion, we re-casted flux networks into reaction-based graphs in which a node represents a reaction and an edge between two reactions represents the mass flow between them mediated by a shared metabolite^49^ (**Methods**, **Figure S3A**). By tracing nutrient-to-biomass paths on reaction-based graphs, we decomposed the mass flow mediated by a specific nutrient into its relative contributions across metabolic subsystem annotations (**Methods**, **Figure S3B**). The resulting decomposition revealed that nutrients are either multifunctional and involved in many subsystems, or specialized and contributing directly to biomass. Whether a nutrient was multifunctional or specialized depended on both the network structure and the environment. For example, arginine was multifunctional in *E. coli* under simple environments but became specialized in the full environment (**Figure S3B**). Regardless of species, the proportion of specialized nutrients increased in the full environment compared to in simple environments (**Figure S3B**). These results suggest that utilizing more nutrients reorganized the flux networks in a similar way across dissimilar stoichiometries, which motivated us to coarse-grain the networks.

To do so, we sampled optimized networks across all species in environments of varying complexity, covering network configurations ranging from the most efficient ones that can be achieved only when all nutrients are available to the less proteome-efficient ones under more limited environments. Then, we summed the reaction-based graphs to obtain a union graph where the weight of an edge quantified whether it is part of a context-specific branch or an emergent backbone shared across contexts. To parse the regulatory information contained in the union graph, we used an algorithm based on random walks on graphs to learn the local and global context of the nodes^50^ – reasoning that local context captures biochemical function while global context captures distance to nutrient utilization and biomass reaction – and finally, clustered the embedded reactions into sectors (**Methods**, **Figure 3A**).

This coarse-graining procedure amounts to an unsupervised identification of metabolic proteome sectors from genomes alone. For the rest of this work, we focused on a particular clustering of 20 sectors that we deemed most suitable as a starting point through experimentation on hyperparameters and manual inspection. We discuss their robustness in the next section. We refer to this clustering as, putatively, the conserved growth predictive (CGP) sectors.

The CGP sectors were interpretable and informative. Biomass reactions for the different species all naturally clustered into one sector, even though their reactant compositions can differ by more than 50% due to, for example, differences in cell envelope compositions between Gram types. Central carbon metabolism formed a prominent backbone and was largely captured by three sectors (**Figure 3B**). Sector 11 contained glycolytic and respiratory reactions that converge on pyruvate; sector 12 contained fermentation modules branching from pyruvate; and sector 3 and 13 contained the TCA cycle. Other sectors were also broadly consistent with metabolic annotations (**Figure S4**), demonstrating that our unsupervised coarse-graining method recovered canonical biochemical organizations.

Importantly, the CGP sectors also captured flux-dependent network features missing from annotations. For example, the reaction PYK, yielding pyruvate, was clustered into the end of sector 11 even though pyruvate is a hub metabolite connected with many sectors. This rational clustering was obtained because the flux-weighted random walks frequently traversed along the glycolysis pathway upstream of PYK, whereas downstream of PYK, traversals were split among multiple pathways, thereby distinguishing the embeddings (**Figure 3C**). The same mechanism was also able to resolve opposite directions of the same metabolic conversion or even the same reaction. The anaplerotic reaction PPC from phosphoenolpyruvate to oxaloacetate clustered with glycolysis into sector 11, whereas the reaction PPCK in the opposite direction clustered into sector 3 along with other gluconeogenesis reactions (**Figure 3B**). In fact, PPCK can proceed in the anaplerotic direction, as was the case for a PYK knockout of *B. subtilis*^51^, and this direction was indeed clustered into sector 11.

With a coarse-graining in hand, we first asked how nutrient contributions to growth depended on network organization, which we quantified via sector fractions *E_i_*, defined for each sector *i* as the sum of fluxes for reactions in that sector divided by the total flux of all reactions. As expected from the decomposition analysis, sector fractions remained largely unperturbed after removing a specialized nutrient from the full environment. For example, dropping out arginine for an *E. coli* model led to substantial changes in only two sectors, one contained multifunctional catabolic reactions and the other contained arginine biosynthesis reactions. The fractions of these two sectors increased and the cost of this compensation was distributed approximately equally across the other sectors, leading to only a small growth loss (**Figure S5A**). By contrast, dropping out a multifunctional nutrient like glycerol 3-phosphate perturbed the entire network, with substantial increases and decreases in fractions across most sectors and a large growth loss (**Figure S5A**). Across species and nutrients, growth loss tended to increase with the difference in sector fractions before and after dropout, and dropouts of specialized (multifunctional) nutrients incurred smaller (larger) sector fraction changes (**Figure S5B**). These results illustrate how metabolic functions can be quantitatively linked to coarse-grained network features.

### Coarse-graining identifies metabolic sectors that correlate with growth across species in expression data

To test the CGP sectors, we analyzed transcriptome and biomass yield data for diverse gut bacterial species grown individually in a complex medium^52^. We calculated sector fractions by taking gene expression as the flux through the corresponding reactions (**Methods**, **Figure 4A**) and found that sector fractions and growth were well correlated: **Figure 4B** shows three such sectors (Spearman’s *ρ* > 0.54 and *p* < 0.02 for all three sectors). These sectors were selected as examples because the mass flows from these sectors to biomass were proteome efficient, as defined in the next section. Across all sectors, the largest correlation coefficient was 0.70 and the mean magnitude was 0.45 (**Figure S6**). While not all sectors are expected to be growth predictors, the mean magnitude nonetheless quantifies the overall performance for this set of sectors. Notably, many sectors had correlations larger than those for the ribosomal fraction, which was itself positively correlated with growth (*ρ* = 0.55, *p* = 0.02; **Figure 4C**). Moreover, the total fraction of all metabolic genes was negatively correlated with growth (*ρ* = −0.57, *p* = 0.01; **Figure 4C**). These latter correlations are reminiscent of bacterial growth laws, suggesting that growth predictors can indeed be conserved. Sector fractions for randomized sectors, obtained by shuffling the assignments from genes to sectors, had significantly smaller mean correlations with growth (*z*-score = 2.4; **Figure 4D**), showing that growth correlations in the CGP sectors were not statistical artifacts.

The status of the CGP sectors as conserved growth predictors is relatively robust also in silico, with respect to model details, and with respect to algorithmic parameters. For simulations across species in the full environment, the CGP sectors were significantly more correlated with growth than shuffled sectors (*z*-score = 6.3; **Figure S7A**). In silico correlations were weaker on average than in vitro (**Figure S6**), but the in silico and in vitro correlations were themselves fairly consistent (*r* = 0.45, *p* = 0.05; **Figure S7B**). We note the same caveats as when interpreting growth correlations with *M*, and thus, comparisons between in silico predictions and in vitro tests are only qualitative.

We next tested the assumption that all species have the same total proteome budget *P*. We randomly assigned species-specific values of *P* from a uniform distribution between the previously assumed budget and half of it (corresponding to a coefficient of variation CV = 0.2). We then performed the same simulations and coarse-graining as for the CGP sectors to obtain variant sectors. These variant sectors also had a mean correlation in vitro that was significantly larger than random shuffles (*z*-score = 2.1; **Figure S7C**). Species-specific-*P* simply rescaled all fluxes: it lowered growth correlations with the number of nutrients utilized *M* in proportion to the CV of *P*, but correlations with *M* remained larger than correlations with input flux even though the latter correlations were now artificially inflated due to network-wide rescaling (**Figure S7D**). Despite this rescaling, the rankings of the in silico growth correlations of the CGP sectors remained largely preserved across instances of species-specific-*P* (**Figure S7E**), supporting the robustness of the coarse-graining.

Algorithmically, coarse-graining performance was not affected by varying the random seeds in the embedding and clustering algorithms (**Figure S7F, G**), or leaving out metabolic models of *E. coli* and species matched to the ones in the experiment (**Figure S7H**). However, a smaller number of clusters (*k* = 10) resulted in growth correlations in vitro that were indistinguishable from random shuffles and biomass reactions clustering together with other reactions (**Figure S7I**), showing that the chosen number of clusters (*k* = 20) was a successful heuristic.

Taken together, the analyses above suggest that coarse-graining metabolic networks successfully identified a set of conserved growth predictors involving expression data without using any gene expression data.

To further test the CGP sectors, we applied the same data analysis framework to transcriptome and growth rate data for *E. coli* across 131 contexts^31^, and found that in this case, the mean correlation of the CGP sectors was not significantly larger than random (*z*-score= 0.08; **Methods**, **Figure S8**). In fact, the total metabolic fraction was also not correlated with growth (*ρ* = −0.16, *p* = 0.07; **Figure S8**). These results indicate that the status of the CGP sectors as conserved growth predictors, like that of *M*, also depended on the ensemble of contexts. The contexts here included various laboratory evolved strains, suggesting that laboratory evolution samples growth regulatory strategies different from the ones captured by the CGP sectors.

By contrast, for an *E. coli* model simulated across wide-ranging nutrient environments, the CGP sectors performed significantly better than random (*z*-score = 3.9; **Figure S8**). We used this setting to test robustness to reaction-specific proteome costs. We simulated a reaction-specific *E. coli* model, for which reactions were assigned specific costs based on enzyme information^53^, across nutrient environments and obtained another set of variant sectors. These sectors performed similarly as the CGP sectors on the simulated *E. coli* dataset (**Figure S9A**). However, unlike species-specific proteome budgets, reaction-specific proteome costs changed inter-reaction relationships: in silico growth correlations for sector 16 switched signs when calculated using the outputs of the reaction-specific model. The proteome fraction of sector 16 decreased by as much as 20% between the reaction-specific and non-specific models, and this decrease was correlated to an increase in the fraction of sector 11 (**Figure S9B**). The switch likely occurred because the glycolytic reactions in sector 11, including GAPD, PGK, PGM, and ENO, had higher enzymatic rates and thus lower proteome costs than average (**Figure S9B**). Except for this and two other outlier sectors, the remaining sectors had similar growth correlations between the reaction-specific and non-specific models. These analyses are consistent with previous studies showing that bacterial growth properties can be explained using simplified cost models^54,55^. To further probe the gap between reaction flux and proteome cost, we tested the CGP sectors in a proteomics dataset comprising *E. coli* across 36 contexts^56^, and found that growth correlations between the transcriptomic and proteomic datasets were largely consistent (**Figure S9C**), in agreement with recent comparisons of transcriptomes and proteomes^57^.

In sum, robustness analyses show that assumptions about the proteome cost model can have specific effects on individual sectors and their growth correlations, which can also depend on the ensemble of species or environments sampled. However, the overall trend between the CGP sectors and growth was relatively robust.

Lastly, we tested the relationship between CGP sectors versus transcription factors and targets. We analyzed iModulons, statistically independent groups of genes extracted via independent component analysis on the same set of *E. coli* transcriptome data as above^31^. Previous work identified 52 metabolism-related iModulons that could be associated by manual curation to known transcription factors. Of these, 33 (63%) were significantly enriched in only one of the CGP sectors (**Figure 4E**), suggesting that CGP sectors could arise through transcription factors. For example, the catabolite repressor/activator Cra iModulon was enriched only in sector 11, which was associated with glycolytic reactions, and indeed contained several genes involved in carbon utilization, including the fruBKA operon for fructose utilization^58^. The Cra iModulon also contained ppsA, which is typically associated with gluconeogenesis^59^, and accordingly, was grouped into sector 3 (**Figure 3B**). This seeming inconsistency, in which a single iModulon, enriched in only one sector, includes genes from two sectors that were associated with opposite metabolic directions, is resolved by the fact that, true to its name, Cra represses fruBKA but activates ppsA^60^ (**Figure 4F**). By contrast, the ArgR iModulon was split into multiple CGP sectors in a biologically plausible manner. It consisted of the arginine biosynthesis genes argA-I and was split into two sectors that separately contained argA-E and argF-I. This split corresponds to a natural branch point in the network at ornithine^61^, and while ArgR represses the entire pathway in *E. coli*, it is reported to not repress argA-E in *Pseudomonas aeruginosa*^62^ (**Figure 4F**).

### Proteome-efficient networks have similar coarse-grained features

In this section, we analyze how coarse-grained networks relate to growth. We began by testing how sector fractions change with the nutrient environment by varying the number of supplied nutrients *N*. Since *N* increased growth by enabling more nutrient specialization, we thought that *N* would drive species to similar sector fraction profiles despite species-specific stoichiometries (**Figure 5A**). However, simulations showed that species differences in sector fractions were just as large in the full environment as in simple environments (*N* = 20; **Figure 5B**), suggesting that sector fractions remained species specific. Instead, sector entropy, or the evenness of reaction fluxes within a sector, tended to decrease with increasing *N*, and species differences in sector entropies were significantly smaller in the full environment (**Methods**, **Figure 5C, D**). These results suggest that environment complexity enables networks to converge onto proteome-efficient subsets of reactions within each sector.

**Figure 5.**
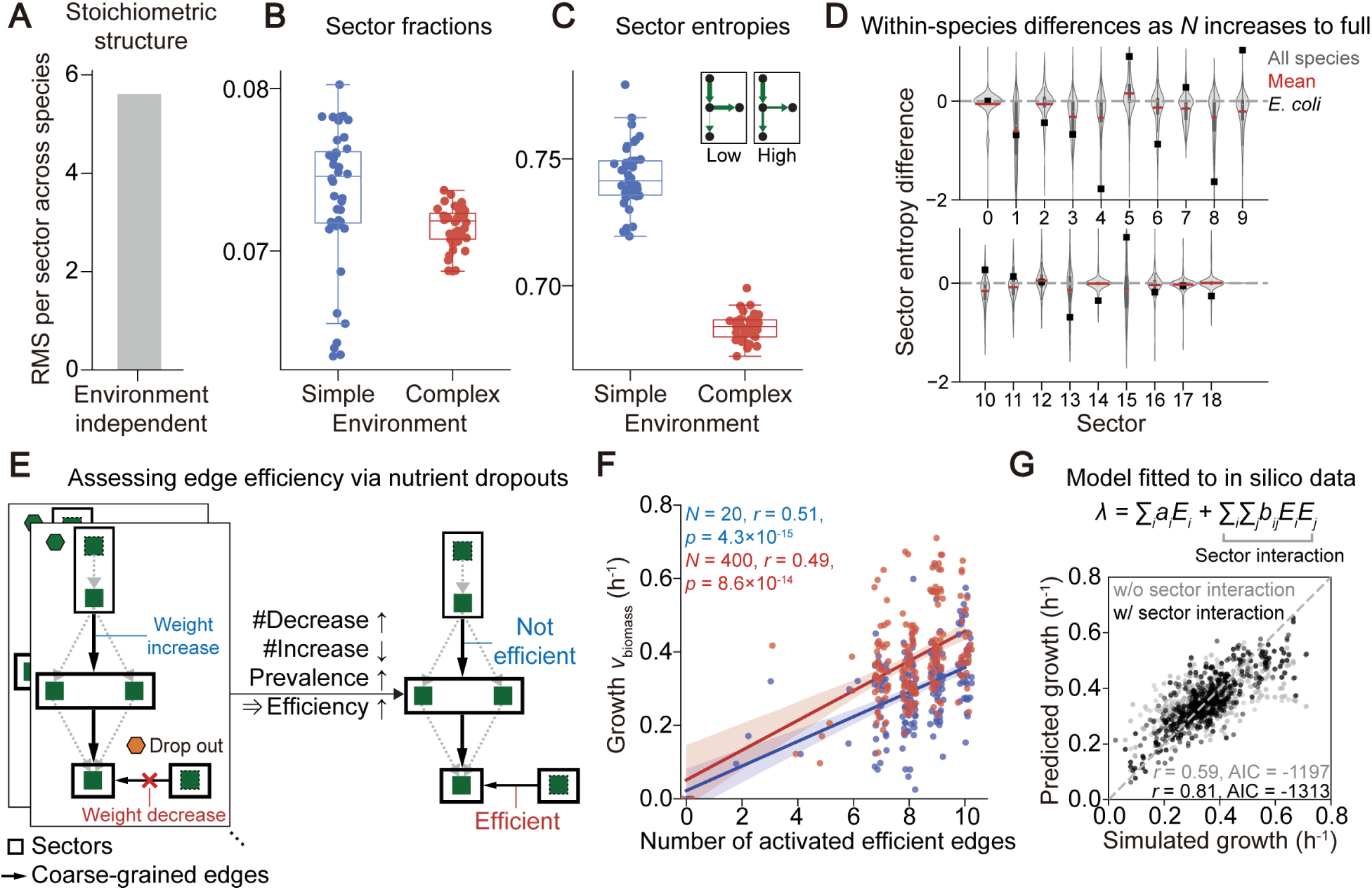
Sector interactions improve growth predictions. A) Species exhibit highly dissimilar stoichiometric structures, as quantified by the root mean square (RMS) of reaction presence-absence divided by 20, the number of CGP sectors (**Methods**). B) Variation in sector fraction profiles across species within a given environment. Shown are RMS per sector values for 40 simple and 40 complex environments. C) Sector entropy profiles are more dissimilar across species than sector fraction profiles, and became more similar across species with increasing environment complexity. Variation in sector entropy profiles as in panel B. D) Sector entropy, for a given species, tended to decrease with increasing environment complexity. Shown are distributions across species of sector entropy differences between simple and complex environments, the mean across species (red), and values for *E. coli* (black). E) Identifying efficient mass flow between sectors via nutrient dropout statistics (**Methods**). Nutrient dropouts induced either decreased or increased edge weights in coarse-grained reaction-based graphs. Edges were deemed efficient if they tended to exhibit decreased weight following dropouts and high frequency before dropouts across species and environments. F) The number of efficient edges activated predicted growth across species and environments. Shown are predictions in simple (*r* = 0.51, *p* = 4e−15; blue) and in complex environments (*r* = 0.49, *p* = 9e−14; red). Solid lines and shaded areas indicate linear fits and 95% confidence intervals. G) Sector interactions contained growth information. Growth rate *λ* was modeled as the sum of contributions from sector fractions *a_i_E_i_* and sector interactions *b_ij_E_i_E_j_* (**Methods**). Models were fitted to simulations across species and across both simple and complex environments. Model complexity was controlled by regularization. The model with sector interaction terms (*r* = 0.81 ± 0.05; black) achieved better predictions at lower Akaike Information Criterion than the model without interactions (*r* = 0.59 ± 0.11; gray).

We hypothesized that sector fractions represent proteome costs, which are fixed per molecule expressed and thus only indirectly dependent on the environment through the expression levels of individual enzymes. By contrast, the growth benefits of individual reactions and coarse-grained sectors should depend on the flux network as a whole. This interpretation is consistent with phenomenological models of bacterial growth in which the flux contribution to growth out of a sector and its enzyme cost are not proportional^24^. To test this hypothesis, we analyzed the coarse-grained edges between sectors, which now represent total mass flow from multiple metabolites, perhaps corresponding to quantities sensed by *E. coli* to regulate ribosomal and other proteome sectors^63^. We assigned an efficiency metric to each edge based on its statistics of activation across nutrient dropouts. Edges that exceeded a threshold mass flow after dropout, suggesting that they were compensatory mass flows activated only under constrained conditions, were deemed less efficient and vice versa (**Methods**, **Figure 5E**). Defining the top ten edges with the highest efficiency metric as efficient, we found that the number of activated efficient edges predicted growth across species and environments, including both complex and simple environments not in the training set of dropouts (*r* = 0.51 and 0.49, *p* = 4e−15 and 9e−14; **Figure 5F**).

This prediction was partially driven by connections to the biomass node, with four of the top ten efficient edges containing biomass precursor metabolites (**Figure S10A**). The top ten efficient edges also contained internal mass flows upstream from the biomass node. In particular, the most efficient internal edge pointed from sector 1 to 12 and was mediated by the conversion of G6P to UDP-glucose, which can then flow into cell wall and hence biomass synthesis (**Figure S4**). That mass flow into sector 12 was efficient is consistent with the mechanism of overflow metabolism, which arises at high growth rates because fermentation – also clustered into sector 12 (**Figure 3B**) – is more proteome efficient than respiration^46^. The identification of efficient edges was robust: their growth correlation was significantly higher than those of randomly selected edges (**Figure S10B**). More generally, growth was better predicted by sector fractions, and at lower Akaike Information Criterion, when pairwise interaction terms between sectors were included (**Methods**, **Figure 5G**). The same model, which was trained only on in silico results, was able to predict the *E. coli* transcriptomic dataset analyzed above (*r* = 0.54, *p* = 2e−11; **Figure S10C**), further suggesting that the CGP sectors captured growth predictive features in addition to sector fractions. Since edge weights are mathematically defined by fluxes from multiple sectors, these results are consistent with the interpretation that growth benefits are encoded by sector interactions.

While a more mechanistic picture remains to be elucidated, these analyses suggest that across diverse metabolic networks, there is an emergent layer of coarse-grained features that govern growth.

## DISCUSSION

We were motivated by several recent experimental findings to search for quantitative trends in bacterial growth that emerge across species and environments. Analyzing genome-scale metabolic models in silico, we found two growth predictors that were simple and conserved: the number of nutrients utilized and the CGP metabolic sectors. Importantly, the sectors were learned from flux networks predicted by proteome-constrained optimization, and thus, can be understood as groups of reactions under similar regulation across contexts. The proteome fractions of the CGP sectors were correlated with growth across tens of species tested in the experimental dataset we analyzed, despite the fact that the sectors were defined without using any gene expression data.

The growth correlations found were moderate, at least compared to the perfectly linear bacterial growth laws. However, they were observed across disparate experiments and relatively robust to various modeling details. We emphasize that the growth correlations are not some universal laws across bacteria. Rather, we interpret them to suggest that, even across quite dissimilar species or nutrient environments, there can exist conserved growth predictors.

Our work has several limitations. The primary limitation is a lack of experimental data. The vast majority of gene expression data for bacteria are obtained for a few model organisms such as *E. coli* and most do not have matching growth measurements. Furthermore, growth measurements are not standardized across studies, with some reporting biomass yield and others reporting growth rate. Nonetheless, trends are emerging from recent studies and the scant data currently available, including the result that the ribosomal and total metabolic fractions were indeed correlated with growth across species in one of the datasets analyzed^52^. Another caveat is that we assumed that all species regulate growth via proteome-constrained optimization, even though other regulatory strategies exist^25,36,64,65^. This assumption is straightforward to relax since FBA and its variants can implement different regulatory algorithms. More broadly, diverse regulatory strategies should lead to more, not fewer, conserved growth predictors: since growth predictors emerge from active regulation rather than static network properties^16,28,64^, the challenge is the inverse problem of finding the combinations of networks, nutrient environments, and regulatory strategies that give rise to growth correlations.

The proposed coarse-graining framework is a key contribution of our work. It finds coarse-grained features of metabolic networks informed by growth regulatory strategies across contexts, and does so from genomes alone. Thus, our framework can be easily extended to obtain coarse-grainings in other contexts, as we did to investigate the robustness of the CGP sectors. Moreover, since our framework defines reaction groups based on metabolic network features, it avoids grouping unrelated confounders or non-metabolic genes together, as can occur when clustering expression data^17,31^. The flexibility of our framework will help elucidate the link between metabolic sectors, network topology, and how different nutrient combinations contribute to overall growth^66,67^. More broadly, conserved growth predictors will facilitate applications involving diverse species, for example by providing simple metrics to interpret metaproteomes as well as key targets to manipulate in situ^68,69^.

## METHODS

### Proteome-constrained growth optimization via flux balance analysis

To simulate growth and metabolic flux networks across species and environments, we applied flux balance analysis (FBA) to genome-scale metabolic models (GEMs)^12^. We selected 205 GEMs from the AGORA2 database of high-quality, manually-curated models of human gut bacteria^37^, corresponding to 205 phylogenetically diverse and representative species (**Table S1**). We simulated these models using FBA with a simple proteome constraint detailed below. To be explicitly clear, the models were always simulated individually, without interactions from other species.

In brief, FBA takes as input a stoichiometric matrix *S* describing the metabolic reactions of the system and solves for the vector *v* of fluxes over the reactions. FBA assumes that the network is at steady state such that the influx to and outflux from a metabolite are equal for every metabolite. The environment is encoded by imposing lower and upper bounds on the fluxes of exchange reactions, which encode the import of nutrients from the external environment into the system. Under these constraints, FBA maximizes the flux through the biomass synthesis reaction, resulting in a linear programming problem.

On top of the basic FBA framework, we implemented a simple proteome constraint. We assumed that each reaction incurred a proteome cost for the cell due to the amount of enzyme required to support the flux. For simplicity, all reactions were assumed to have the same cost coefficient^54,70^. The resulting optimization problem is

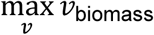

subject to

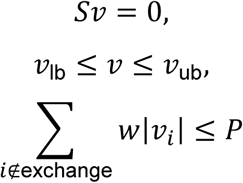

Here, *v*_biomass_ denotes the flux over the biomass reaction which is also the growth rate *λ*. *v*_lb_ and *v*_ub_ are the lower and upper bounds of the flux vector *v*, with units of mmol*h^-1^*gDW^-1^ ^12^. *w* is the cost coefficient and was set to 2.6 × 10^-4^ g*mmol^-1^*h, representing the average value across all enzymes in *E. coli* of the ratio of the molecular weight of an enzyme and its kinetic efficiency^71^. *P* is the total proteome budget and was set to 0.30 g*gDW^-1^, corresponding to an upper bound of 30% on the fraction of cellular dry mass dedicated to metabolism^16,72^. The proteome constraint was applied to all reactions except the biomass reaction and exchange reactions for the base medium (below).

The environment was specified by the set of supplied nutrients. All environments included a base medium containing inorganic ions and gases (Mn^2+^, Cu^2+^, HPO_4_^2-^, Co^2+^, H^+^, Mg^2+^, CO_2_, O_2_, Cl^-^, Zn^2+^, SO_4_^2-^, Fe^3+^, H_2_O, K^+^, NH_4_^+^, Ca^2+^, Na^+^, Fe^2+^), which were assumed to be present in excess, with the lower bound of the corresponding exchange reactions set to −1,000 mmol*h^-1^*gDW^-1^. Additional nutrients were supplied depending on the environment, with the lower bound of their exchange reactions set to −20 mmol*h^-1^*gDW^-1^ to ensure saturating conditions^12^. To bring the resulting growth rates into a range comparable to that in experiments, all flux values shown in the figures were rescaled by dividing by a constant factor of 5.

All simulations were performed by pFBA^73^ using COBRApy (version 0.29.0)^74^, with optimization solved using Gurobi^75^.

### Universal model

For various downstream purposes, we constructed a “universal” model by aggregating all metabolic reactions from the 205 selected GEMs. Thermodynamically infeasible energy-generating cycles – defined here as those that can produce energy metabolites (i.e., ATP, CTP, GTP, ITP, NADH, NADPH, flavin adenine dinucleotide, flavin mononucleotide, menaquinol, 2-demethylmenaquinol, acetyl-CoA, L-glutamate, and proton) without nutrient uptake – were removed by minimally deleting reactions via GlobalFit^48^. The resulting model comprised 7,425 reactions and 4,529 metabolites.

### Randomly generated metabolic networks

We sought to generate thermodynamically feasible but evolutionarily implausible network structures. To do so, we performed a random walk in which each step consisted of deleting one and adding one reaction. The reaction to be deleted was randomly selected from the existing reactions. The reaction to be added was randomly selected from all other reactions in the universal model. Only intracellular reactions were added or deleted; transport, exchange, and sink reactions were not added or deleted. If a step decreased the growth rate below 0.002 h^-1^, then it was rejected. This procedure was repeated for 5,000 steps for each random network. We generated a collection of 151 random networks, all initialized from the “*E. coli* K-12 substr MG1655” model.

### Reaction-based graphs

In reaction-based graphs, nodes represent reactions and directed edges between nodes represent mass flow between reactions mediated by shared metabolites. Reaction-based graphs were constructed from optimized flux solutions following Ref. ^49^. In brief, given a stoichiometric matrix *S* and a flux vector *v*, a weighted adjacency matrix between reactions was calculated. Currency metabolites (i.e., ATP/ADP, NAD(H)/NADP(H), phosphate, protons, and water) obfuscated the networks and were omitted by setting their reaction coefficients in *S* to zero, *S_μi_* = 0 for currency metabolite *μ* and all reactions *i*. Reactions were unfolded into forward and backward components; thus, both the flux vector and the stoichiometric matrix were unfolded. The reaction-based adjacency matrix was expressed as an expression of the unfolded *S* and *v*, such that the weight of an edge from reaction *i* to *j* was the sum over all metabolites *μ* of the flow of *μ* produced by *i* multiplied by the flow of *μ* consumed by *j* and divided by the total consumption flow of *μ*. Edges with weight less than 0.002 were excluded to reduce computing time.

### Nutrient function decomposition on reaction-based graphs

To attribute how much a nutrient contributes to various subsystems, we traced nutrient-to-biomass paths on reaction-based graphs and calculated a simple heuristic decomposition as follows. First, each edge was assigned the metabolic subsystem given by model annotations for the reaction targeted by the edge. Then, for each nutrient *μ*, all paths from the nutrient uptake reaction to the biomass reaction, without repeated nodes and less than a maximum path length of 30, were enumerated using a depth-first search algorithm. The contribution of nutrient *μ* to subsystem *y* was calculated as

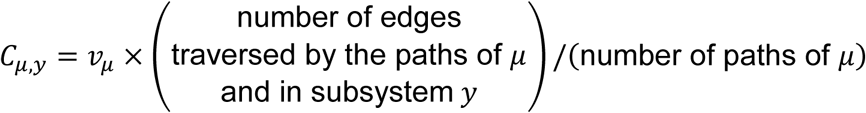

where *v_μ_* denotes the uptake flux of nutrient *μ*. Transport, exchange, and demand reactions were not tallied. The relative contribution 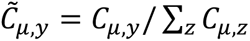 was used to classify nutrient *μ* as specialized if 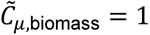 or as multifunctional otherwise.

### Coarse-graining metabolic networks

We sought to obtain a coarse-grained representation of the metabolic network organizations that emerge when diverse species use a shared growth regulatory strategy across environments. To do so, the following simulations were collected: (i) every species in the full environment; (ii) the universal model supplied with the same sets of nutrients utilized by each species in the full environment, representing the most proteome-efficient metabolic states; and (iii) every species in forty randomly selected simple environments, sampling various context-specific pathways and less efficient metabolic states. Reaction-based graphs were obtained and summed to form the union graph. On this union graph, node embeddings were learned using node2vec^50^. node2vec maps nodes to vectors in a low-dimensional space that maximizes the likelihood of preserving the network neighborhoods of nodes, as defined for a node by the collection of nodes reachable from it via random walks. node2vec was applied to the union graph with hyperparameters dimensions=64, and p=1, q=1. The parameters p and q control the probabilities of random walks to backtrack to the previous node or to explore outward nodes, which were set to the values here to implement an unbiased random walk. A fixed random seed was used for reproducibility. The resulting node embeddings were clustered using k-means^76^. Thus, each reaction was assigned to one cluster, or proteome sector, thereby defining the CGP sectors. Finally, for each reaction-based graph, a coarse-grained graph was constructed where the nodes represent the CGP sectors and the edges represent coarse-grained mass flows between sectors. Edge weights were defined as the average weight in the original reaction-based graph of all directed edges between reactions belonging to the corresponding sectors.

### Analysis of expression data

#### Dataset for cross-species analysis

Data were obtained from Ricaurte et al.^52^ for 18 gut bacterial strains grown individually in the complex medium modified Gifu Anaerobic Medium. Final OD was measured at 48 h after growth and taken to be the biomass yield. Transcriptomes were obtained for samples collected during exponential growth.

#### Dataset for cross-environment analysis

Data were obtained from Sastry et al.^31^, which compiled the PRECISE (Precision RNA-seq Expression Compendium for Independent Signal Exploration) dataset. PRECISE contains RNA-seq expression profiles across 154 experimental conditions for *E. coli*, all generated in a single laboratory using a standardized protocol. We restricted our analysis to conditions with growth rates ranging from 0.2 to 1.3 h^-1^, at pH 7, and in the absence of antibiotics, resulting in 131 conditions.

#### Mapping gene expression to reaction cost and sector fraction

Reactions were identified by SEED identifiers in the GEMs that we used^77^, while in the expression data that we analyzed, genes were identified by their sequences. The rest of the genomes were also available. We therefore assigned genes to SEED identifiers by using ModelSEEDpy to reconstruct GEMs from genomes in an automated manner. The resulting GEMs were only used to obtain gene-to-reaction mappings and were not used for further simulations. Functional annotations were also obtained using RAST during model reconstruction^78^.

To calculate sector fractions given gene expression data, each reaction was assigned a proteome cost proportional to the expression level of its corresponding gene. If a gene maps to multiple reactions, then its expression level was distributed equally to the mapped reactions. If a reaction was mapped to by multiple genes, the contributions of each gene were summed. Reaction costs were normalized by the total expression level across genes. Genes not mapped to reactions were omitted. Finally, sector fractions were obtained for each sector by summing the (normalized) proteome costs of the reactions in that sector.

### Comparison between iModulons and CGP sectors

iModulons are independently regulated gene modules identified in *E. coli* by applying independent component analysis to the PRECISE transcriptomic compendium, as previously described^31^. To assess whether CGP sectors capture transcriptional regulatory structure, we tested whether each TF-associated iModulon was significantly enriched in a single CGP sector. To remove spurious associations, iModulons whose dominant sector contained only one gene were excluded, and for a given iModulon, sectors containing fewer than three genes were not considered. Enrichment of each iModulon across sectors was evaluated using Fisher’s exact test with Benjamini–Hochberg correction^79^, with FDR < 0.05 as the significance threshold.

### Cross-species variation in sector-level flux organization

To quantify how metabolic organization varies across species in a given environment, we calculated for several metrics the root mean square (RMS) of the distances between the profile of each species to the average profile across species. RMS per sector was defined as the RMS value divided by 20, the number of CGP sectors.

We considered three metrics: stoichiometric structure, sector fractions, and sector entropies. The stoichiometric structure of a species was represented by a binary vector indicating, in the space of all reactions in the universal model, the presence or absence of reactions in the species. The sector fraction *E_i_* of sector *i* was defined as

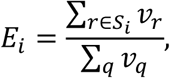

where *S_i_* denotes the set of reactions in sector *i* and *v_r_* the flux of reaction *r*. The sector entropy *H_i_* was defined as

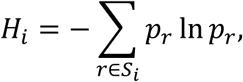

where 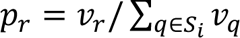 is the normalized flux of reaction *r* within sector *i*. Reactions with zero flux were omitted, and sectors with zero total flux were assigned zero entropy.

### Identification of efficient sector interactions by nutrient dropout

To assess the efficiency of sector interactions, we calculated an efficiency metric for each directed edge between sectors in the coarse-grained reaction-based graph as follows. Coarse-grained reaction-based graphs were constructed for all species grown in the full environment and all nutrient dropouts across species. Dropouts of auxotrophic nutrients that resulted in no growth were excluded. Edge weights were normalized by growth rate and used to assign three properties for each edge. Prevalence was defined as the number of species in which the edge weight was nonzero in the full environment. #Decrease and #Increase were the numbers of dropouts in which the edge weight decreased or increased, respectively, more than a baseline threshold of 0.003 compared to the full environment. Edge efficiency was then calculated as Prevalence × (#Decrease − #Increase + 20), where the constant 20 was set to increase the relative weight of Prevalence.

### Predicting growth from sector fractions

To test the extent to which sector interactions determine growth, we modeled growth rate *λ* as a function of sector fractions *E_i_*,

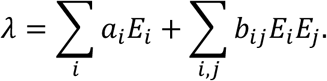

For simplicity, the interaction coefficients *b_ij_* were constrained to be negative, zero, or positive with interaction strength *b*_0_, i.e., *b_ij_* ∈ {−*b*_0_, 0, *b*_0_}. The fitting was done on simulation data for all 205 species grown in four environments of varying complexity (*N* = 20, 60, 400 and 423) under L2 regularization. When interaction terms were included, *b*_0_ was set to 30. The resulting mixed-integer quadratic program was solved using Gurobi^75^. Model performance was evaluated using the Akaike information criterion, AIC = *n* ln(RSS/*n*) + 2*k*, where RSS is the residual sum of squares, *n* is the number of samples, and *k* is the number of fitted parameters that are nonzero.

## ACKNOWLEDGMENTS

We thank members of the Ho lab for helpful discussions. This work was funded by the National Key R&D Program of China (grant no. 2024YFA0920200), the Fundamental and Interdisciplinary Disciplines Breakthrough Plan of the Ministry of Education of China (grant no. JYB2025XDXM502), the National Natural Science Foundation of China (grant no. 32571871), and the Westlake Education Foundation (Competitive Research Funding Program of Center for Synthetic Biology and Integrated Bioengineering, WU2022A002).

## Author contributions

A. Z. and P.-Y. H. designed the research, performed the research, and wrote the manuscript.

## Data availability

The experimental datasets analyzed in this study were obtained from previously published studies and are available in https://github.com/complicatedZ/CGP/tree/main/Expression_datasets. The selected AGORA2 metabolic models used in this study are available at https://github.com/complicatedZ/CGP/tree/main/AGORA2_models. The original sources of the datasets and models are cited in the manuscript.

## Code availability

All code is available at https://github.com/complicatedZ/CGP.

## SUPPLEMENT FIGURES

**Supplement Figure 1.**
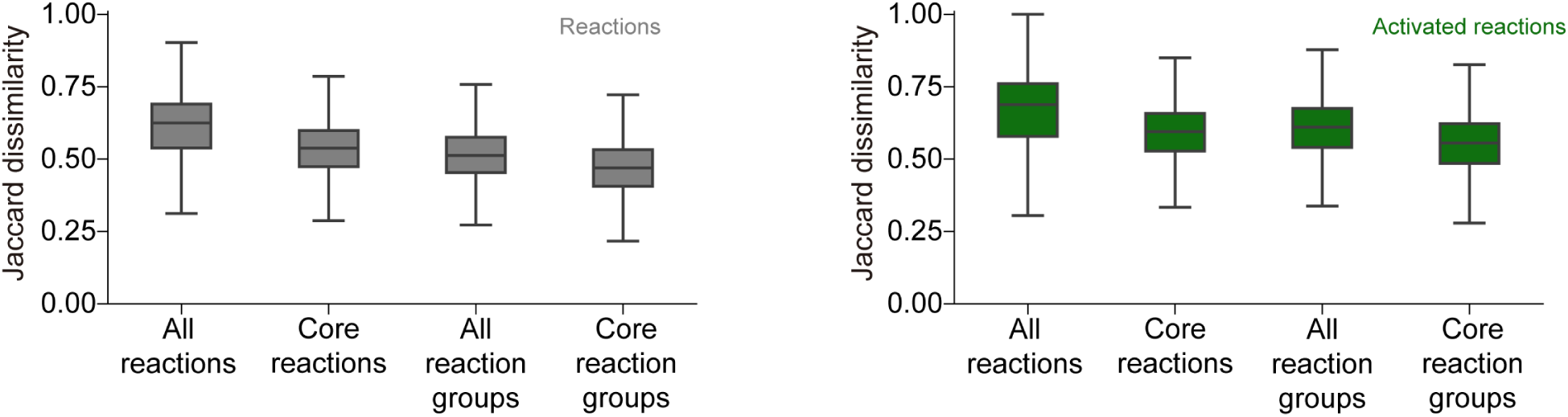
Dissimilarity of metabolic networks. Shown are distributions of the Jaccard dissimilarity between network pairs. “Core reactions” indicate those that belong to the following six KEGG categories, representing central metabolic pathways: glycolysis/gluconeogenesis, the pentose phosphate pathway, the citrate cycle, glyoxylate and dicarboxylate metabolism, pyruvate metabolism, and oxidative phosphorylation. “Reaction groups” indicate groups of similar reactions as defined by the “metabolic pathways” map in iPath3^80^; these groups can include, for example, reactions with slight differences in reactants or reactions from multiple KOs. “Activated reactions” denote those that carry flux in the FBA output in the full nutrient environment.

**Supplement Figure 2.**
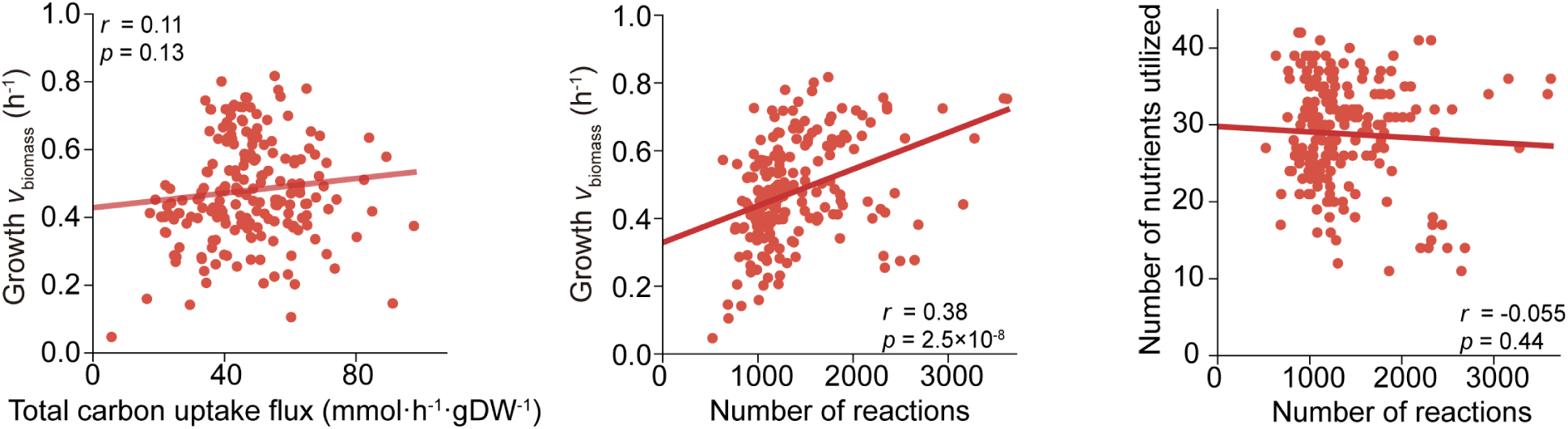
Relationships between growth and network properties in silico across species. Simulations are the same ones as in Figure 2C.

**Supplement Figure 3.**
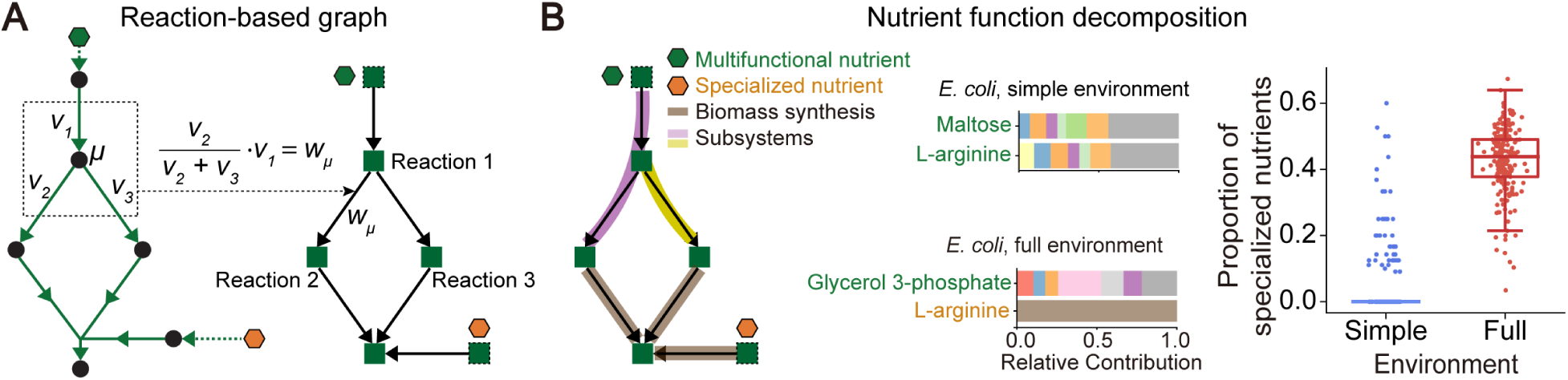
Reaction-based graphs and functional decomposition. A) Re-casting optimized flux solutions into reaction-based graphs. An edge represents the mass flow between two reactions, with weight proportional to the flux of the incoming reaction divided by the total flux across all reactions that consume the mediating metabolite (**Methods**). B) Nutrients became more specialized as environment complexity increased. Left: The function of a nutrient was decomposed by tracing simple paths from the nutrient to the biomass (**Methods**). Based on the decomposition, nutrients were classified as either multifunctional and contributing flux to multiple metabolic subsystems, or specialized and contributing directly to biomass synthesis. Middle: Examples from *E. coli*. In a simple environment, arginine was multifunctional, whereas it became specialized in the full environment. Subsystems with relative contributions below 0.05 were aggregated into a single category (gray). Right: Across species, the proportion of utilized nutrients that were specialized was higher in the full environment than in a simple environment (*N* = 20 nutrients).

**Supplement Figure 4.**
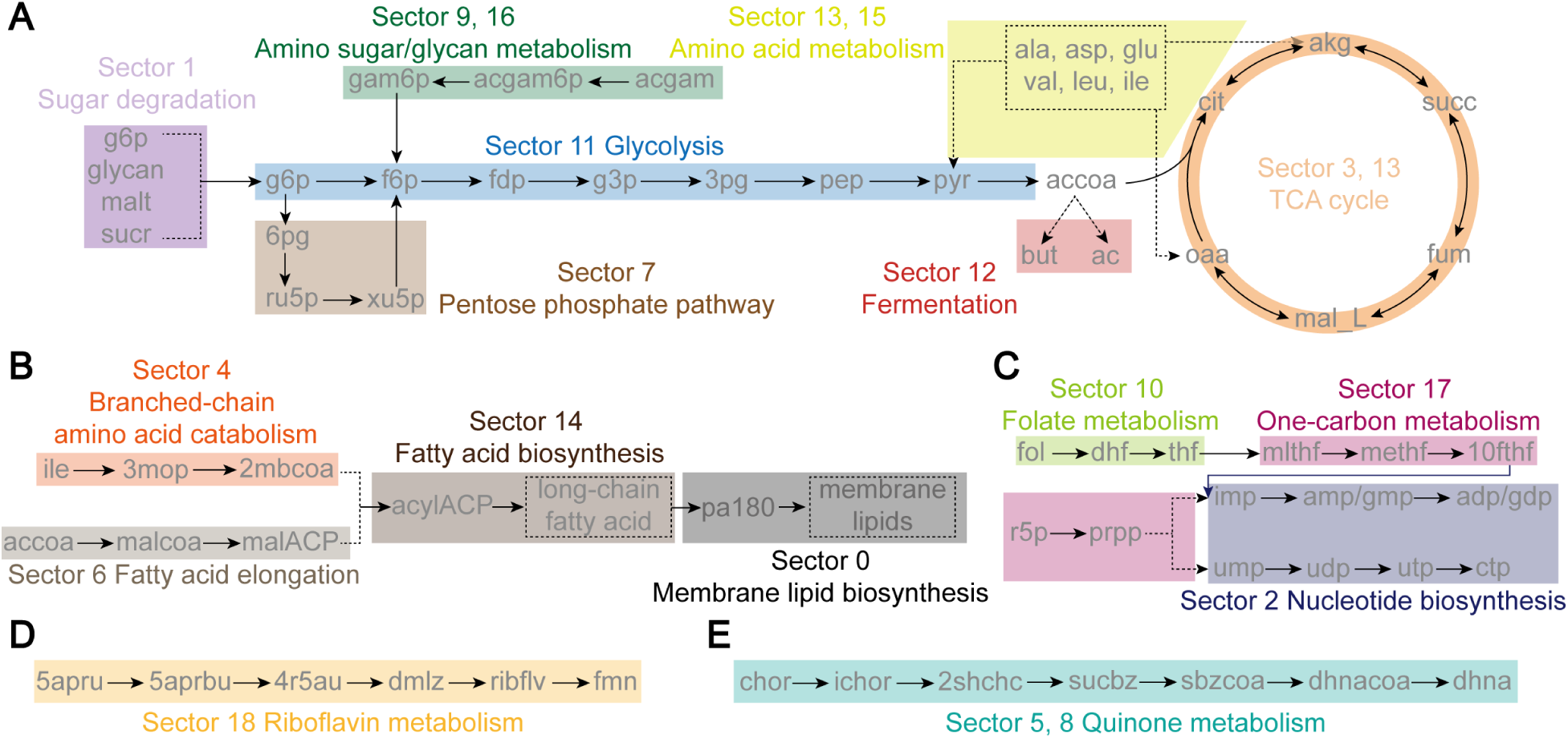
The CGP sectors capture functional metabolic organization. A) Central carbon metabolism, including glycolysis (sector 11), the pentose phosphate pathway (sector 7), the TCA cycle (sectors 3 and 13), fermentation (sector 12), amino sugar/glycan metabolism (sector 9 and 16), and amino acid metabolism (sectors 13 and 15). Colors indicate reaction–sector assignments. Note that sectors contain reactions across many subsystems, and the reactions depicted are only some of the salient features in the sectors. Solid arrows indicate transformations that can be carried out by a single reaction. Dashed arrows or boxes indicate compressed multi-step transformations or downstream endpoints. B) Lipid metabolism, including branched-chain amino acid catabolism (sector 4), fatty acid elongation (sector 6), fatty acid biosynthesis (sector 14), and membrane lipid biosynthesis (sector 0). C) Nucleotide metabolism, showing connections between folate metabolism (sector 10), one-carbon metabolism (sector 17), and nucleotide biosynthesis (sector 2). D) Riboflavin metabolism (sector 18). E) Quinone metabolism (Sector 5 and 8).

**Supplement Figure 5.**
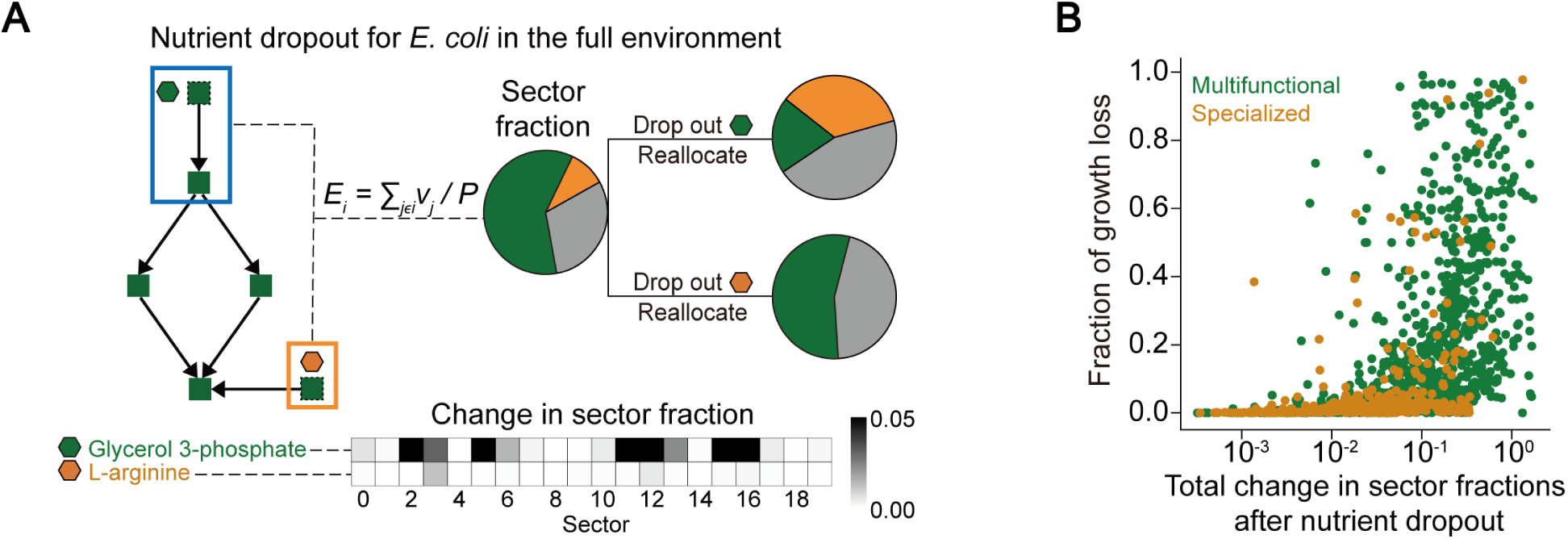
Nutrient dropouts versus sector fractions. A) Nutrient dropout induced reallocation of sector fractions according to nutrient function. Sector fraction is defined as *E_i_* = ∑*_j_*_∈*i*_ *v_j_* /*P*, where the sum is over all reactions *j* in sector *i*. For *E. coli* in the full environment, dropouts of glycerol 3-phosphate, a multifunctional nutrient, and L-arginine, a specialized nutrient, led to different patterns of sector fraction reallocations. B) Dropouts of multifunctional nutrients, compared to dropouts of specialized nutrients, led to larger fractions of growth loss and larger total change in sector fractions after dropout. Across all dropouts, excluding dropouts of auxotrophic nutrients that led to no growth, larger growth loss tended to be accompanied by larger sector reallocations.

**Supplement Figure 6.**
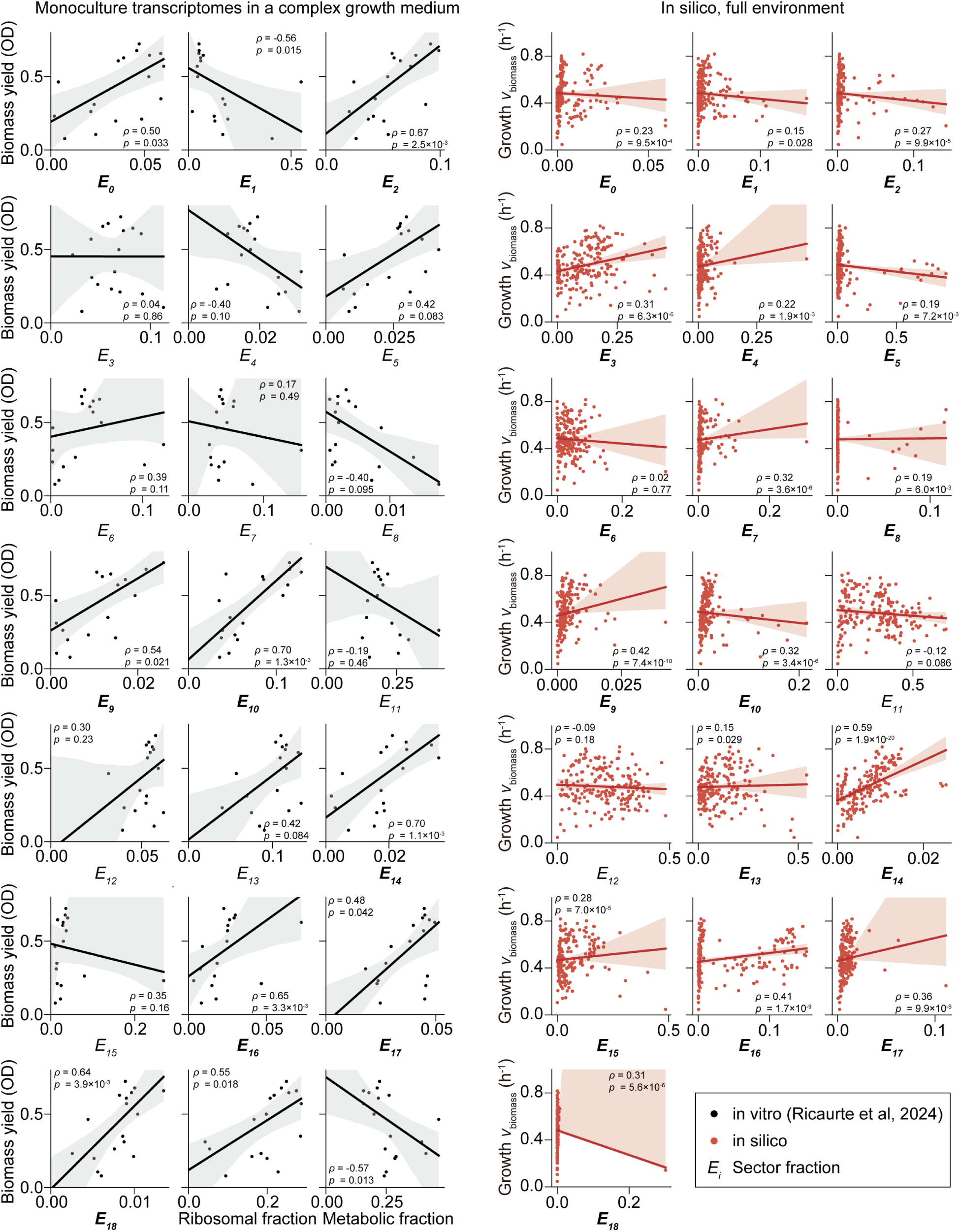
Growth correlations for the CGP sectors across species. Relationship between growth and sector fraction, as in Figure 4B, for all sectors. Solid lines and shaded areas indicate linear fits and 95% confidence intervals. Bold sector label on the x-axis indicates significant correlation.

**Supplement Figure 7.**
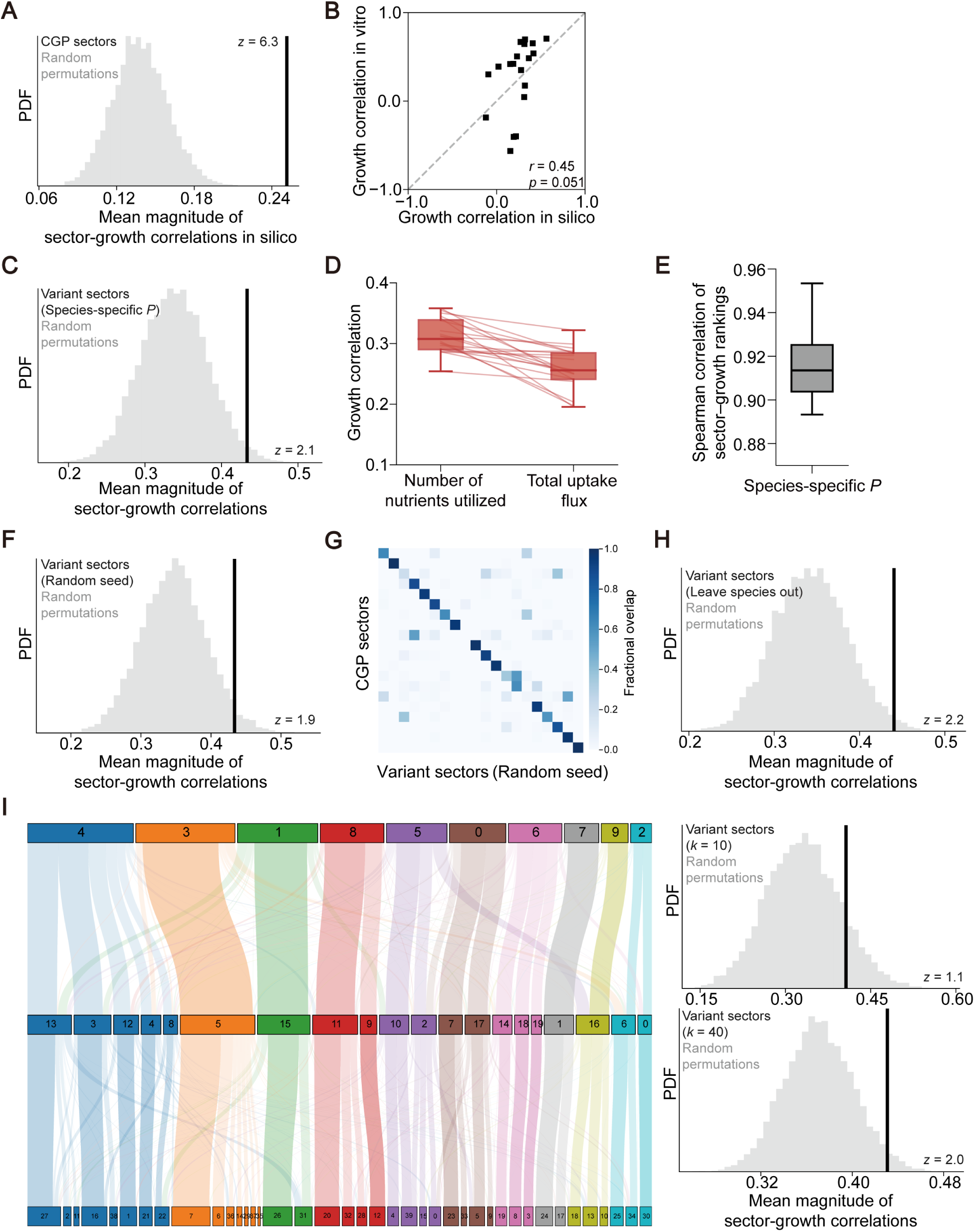
Robustness analyses of the CGP sectors. A) Overall performance, as in Figure 4D, for in silico data. B) In vitro and in silico growth correlations were fairly consistent (*r* = 0.45, *p* = 0.05). Shown are the 19 CGP sectors excluding the biomass sector. C) Overall performance in vitro for a set of variant sectors identified by repeating the coarse-graining procedure in Figure 3A, but for simulations using species-specific total proteome budgets *P*. D) Comparison of growth correlations with the number of nutrients utilized and total uptake flux across 20 instances of species-specific-*P* distributions. Lines connect results from the same instance. E) Shown are Spearman correlations between the sector-growth correlations calculated from species-specific-*P* simulations versus those from the original uniform-*P* simulations. F) Overall performance in vitro for variant sectors identified by repeating the coarse-graining procedure using different random seeds for embedding. G) Fraction of overlapping reactions between the original CGP sectors and variant sectors identified using a different random seed for clustering. H) Overall performance in vitro for variant sectors identified by repeating the coarse-graining procedure, but excluding simulations of species in the experimental datasets analyzed. I) Left: Sankey diagram for the CGP sectors, which was obtained by clustering to *k* = 20 sectors, and two sets of variant sectors identified by clustering to *k* = 10 and 40. Right: overall performance in vitro for variant sectors identified at different *k*.

**Supplement Figure 8.**
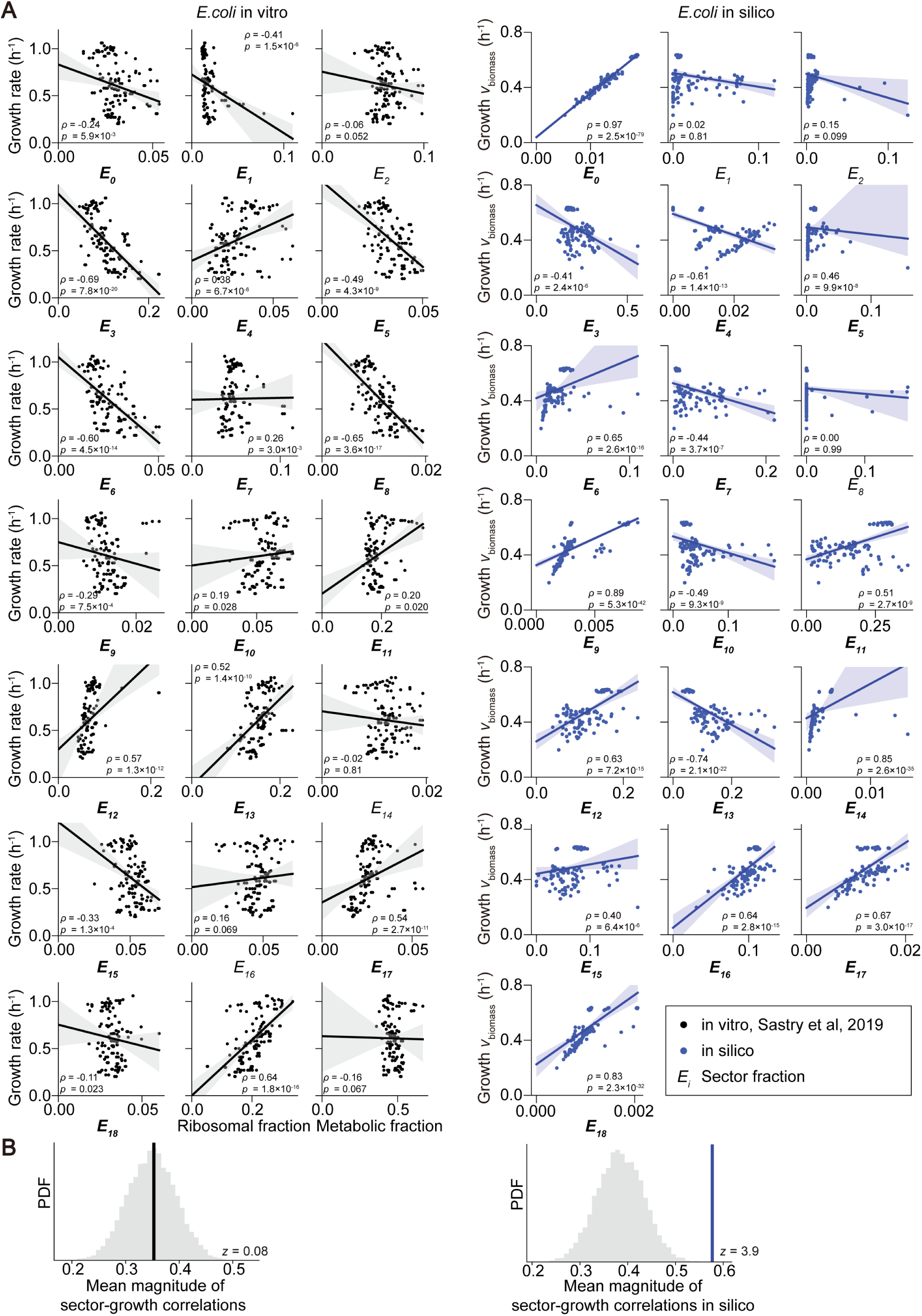
**Growth correlations for the CGP sectors across *E. coli* strains and nutrient environments.**A) Same as **Figure S6**, but for transcriptome and growth rate data from Ref.^31^. The data were obtained for various *E. coli* strains and nutrient environments, amounting to 131 contexts. Solid lines and shaded areas indicate linear fits and 95% confidence intervals. Bold sector label on the x-axis indicates significant correlation. B) Overall performance of the CGP sectors for the in vitro (left) and in silico (right) datasets shown in panel A.

**Supplement Figure 9.**
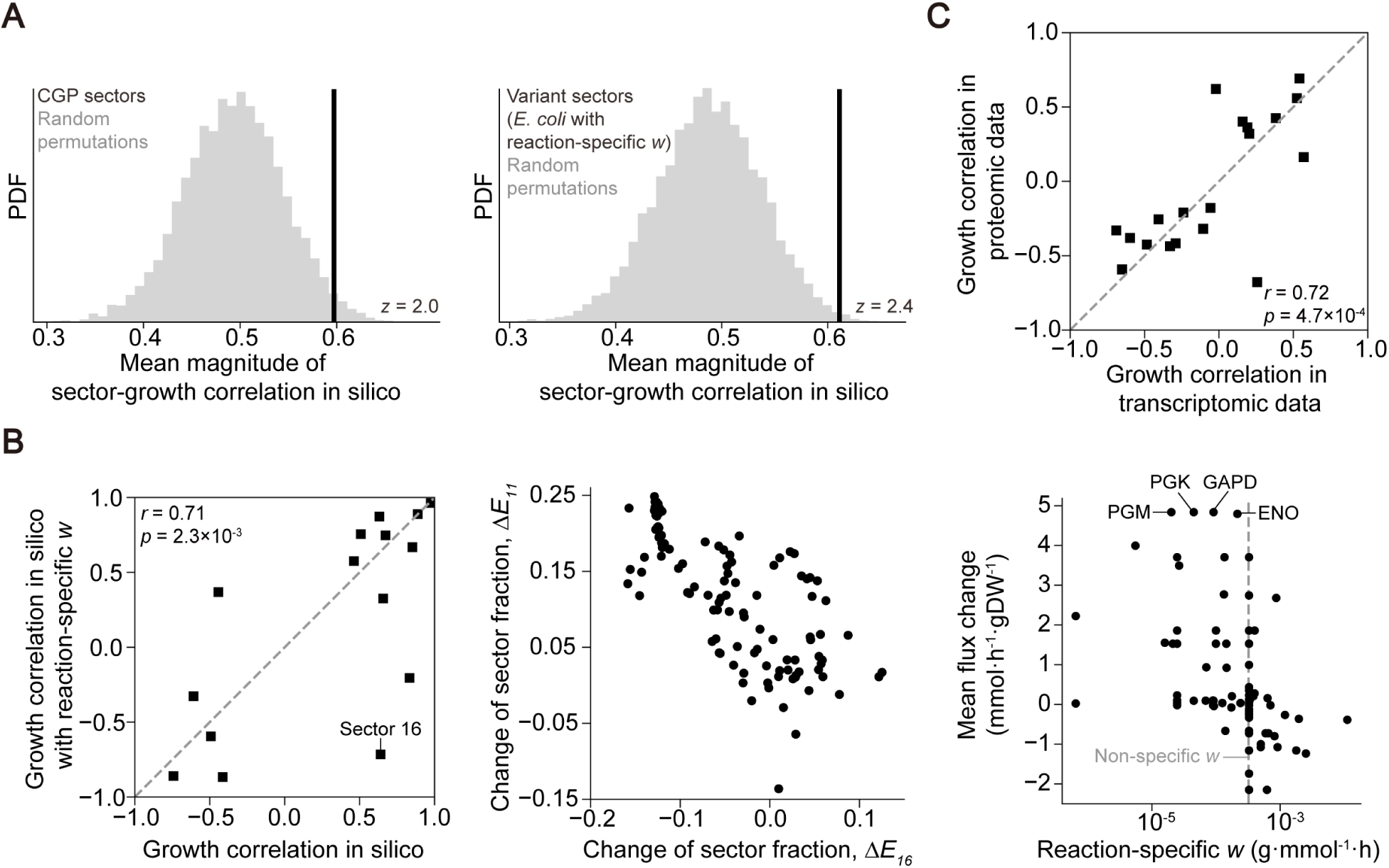
Robustness analyses of the CGP sectors to reaction-specific proteome costs. A) Left: overall performance of the CGP sectors for an in silico dataset generated by simulating an *E. coli* model with reaction-specific weights *w* for proteome costs, defined based on enzyme information of molecular weights and kinetic rates^53^, across 131 wide-ranging nutrient environments. Right: overall performance of variant sectors identified by coarse-graining the same set of simulations of the reaction-specific model. B) Left: growth correlations of the CGP sectors, for simulations using the reaction-specific model and for the original simulations using the non-specific model. CGP sectors with *p* > 0.05 in both settings were excluded. Middle: Relationship between changes in sector fractions for sectors 11 and 16; each data point shows, for a given nutrient environment, the change in sector fraction between the reaction-specific and non-specific models. Right: Flux change, averaged over the simulated nutrient environments, versus reaction-specific *w*; each data point shows a reaction in sector 11. C) Growth correlations in a proteomic dataset comprising *E. coli* in 36 conditions, versus those in the transcriptome dataset analyzed in **Figure S8**; each data point represents one CGP sector.

**Supplement Figure 10.**
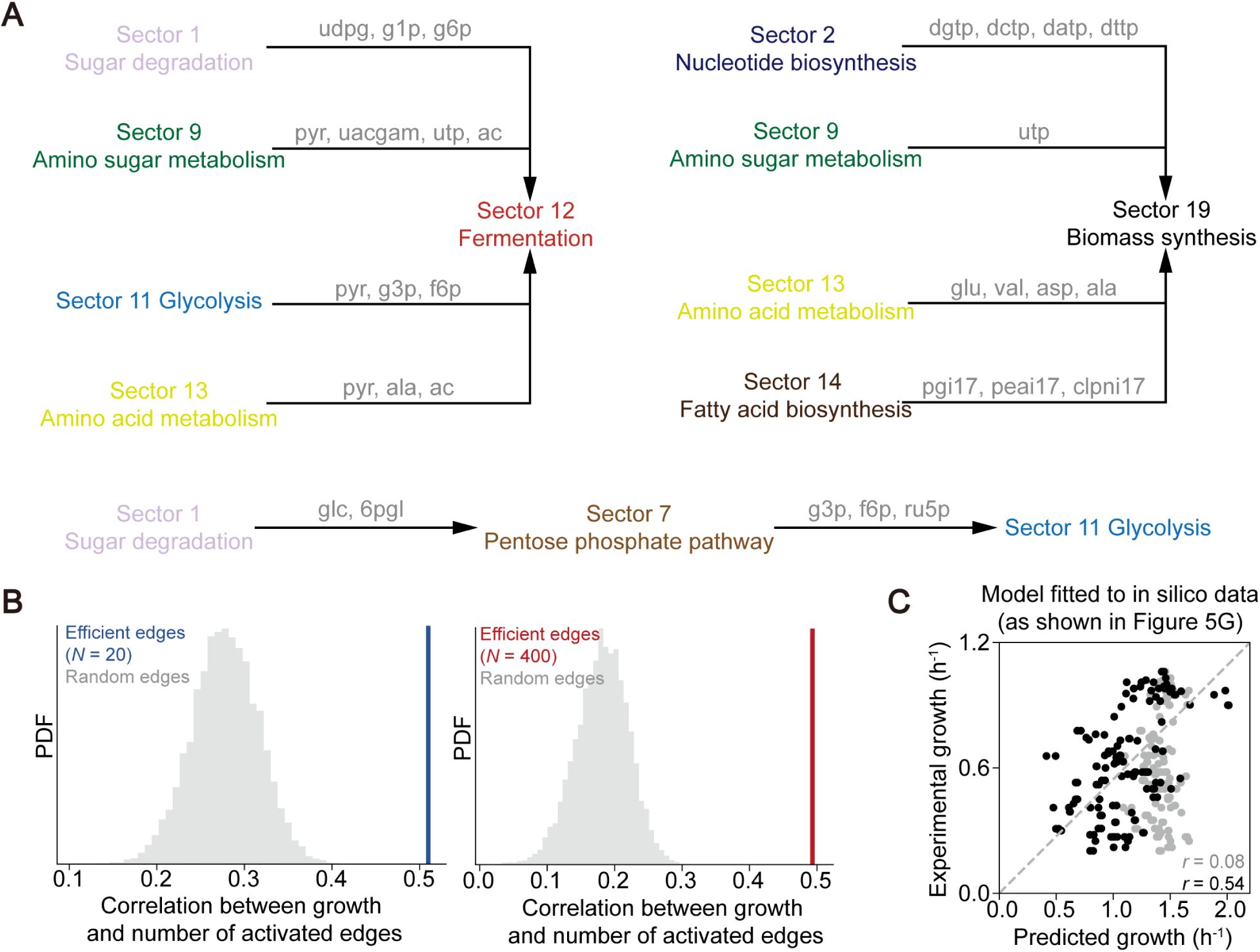
Efficient edges and sector interactions. A) Shown are the top ten efficient edges between the CGP sectors. Representative contributing metabolites are labeled along each edge. Colors denote sector identities as in **Figure S4**. B) Shown is the correlation between growth and the number of activated edges among the top 10 efficient edges (as in Figure 5F), compared to that for sets of 10 randomly selected edges. Correlations are shown separately for simulations in simple (*N* = 20 nutrients supplied) and complex (*N* = 400) environments. C) Growth rates in vitro were predicted by sector fractions using a simple linear model inferred from in silico results (as shown in Figure 5G). Shown are predictions without (*r* = 0.08, *p* = 0.32; gray) and with sector interaction terms (*r* = 0.54, *p* = 2e−11; black).

**Table S1.** List of genome-scale metabolic models analyzed.

| Genus | Species | Model Name |
| --- | --- | --- |
| *Acidaminococcus* | *fermentans* | Acidaminococcus_fermentans_DSM_20731 |
| *Acinetobacter* | *baumannii* | Acinetobacter_baumannii_ATCC_19606 |
| *Acinetobacter* | *haemolyticus* | Acinetobacter_haemolyticus_ATCC_19194 |
| *Adlercreutzia* | *equolifaciens* | Adlercreutzia_equolifaciens_DSM_19450 |
| *Akkermansia* | *muciniphila* | Akkermansia_muciniphila_ATCC_BAA_835 |
| *Alistipes* | *onderdonkii* | Alistipes_onderdonkii_DSM_19147 |
| *Bacillus* | *subtilis* | Bacillus_subtilis_B_1 |
| *Bacteroides* | *caccae* | Bacteroides_caccae_ATCC_43185 |
| *Bacteroides* | *cellulosilyticus* | Bacteroides_cellulosilyticus_DSM_14838 |
| *Bacteroides* | *eggerthii* | Bacteroides_eggerthii_DSM_20697 |
| *Bacteroides* | *fragilis* | Bacteroides_fragilis_NCTC_9343 |
| *Bacteroides* | *stercoris* | Bacteroides_stercoris_ATCC_43183 |
| *Bacteroides* | *thetaiotaomicron* | Bacteroides_thetaiotaomicron_VPI_5482 |
| *Bacteroides* | *uniformis* | Bacteroides_uniformis_ATCC_8492 |
| *Bacteroides* | *vulgatus* | Bacteroides_vulgatus_ATCC_8482 |
| *Bifidobacterium* | *adolescentis* | Bifidobacterium_adolescentis_ATCC_15703 |
| *Bifidobacterium* | *breve* | Bifidobacterium_breve_DSM_20213 |
| *Bifidobacterium* | *longum* | Bifidobacterium_longum_NCC2705 |
| *Bilophila* | *wadsworthia* | Bilophila_wadsworthia_ATCC_49260 |
| *Blautia* | *hydrogenotrophica* | Blautia_hydrogenotrophica_DSM_10507 |
| *Blautia* | *producta* | Blautia_producta_DSM_2950 |
| *Butyrivibrio* | *virosa* | Butyrivibrio_virosa_DSM_23226 |
| *Campylobacter* | *jejuni* | Campylobacter_jejuni_NCTC11351 |
| *Citrobacter* | *freundii* | Citrobacter_freundii_ATCC_8090 |
| *Clostridioides* | *difficile* | Clostridioides_difficile_ATCC_9689 |
| *Clostridium* | *bolteae* | Clostridium_bolteae_ATCC_BAA_613 |
| *Clostridium* | *clostridioforme* | Clostridium_clostridioforme_YL32 |
| *Clostridium* | *hathewayi* | Clostridium_hathewayi_DSM_13479 |
| *Clostridium* | *hylemonae* | Clostridium_hylemonae_DSM_15053 |
| *Clostridium* | *scindens* | Clostridium_scindens_ATCC_35704 |
| *Clostridium* | *symbiosum* | Clostridium_symbiosum_ATCC_14940 |
| *Colibacter* | *massiliensis* | Colibacter_massiliensis_DSM_103304 |
| *Collinsella* | *aerofaciens* | Collinsella_aerofaciens_ATCC_25986 |
| *Coprococcus* | *comes* | Coprococcus_comes_ATCC_27758 |
| *Corynebacterium* | *accolens* | Corynebacterium_accolens_ATCC_49725 |
| *Cupriavidus* | *metallidurans* | Cupriavidus_metalldurans_CH34 |
| *Cutibacterium* | *acnes* | Cutibacterium_acnes_ATCC_11828 |
| *Desulfovibrio* | *piger* | Desulfovibrio_piger_ATCC_29098 |
| *Dialister* | *invisus* | Dialister_invisus_DSM_15470 |
| *Dorea* | *formicigenerans* | Dorea_formicigenerans_ATCC_27755 |
| *Eggerthella* | *lenta* | Eggerthella_lenta_DSM_2243 |
| *Enterobacter* | *cloacae* | Enterobacter_cloacae_subsp_cloacae_ATCC_13047 |
| *Enterococcus* | *dispar* | Enterococcus_dispar_ATCC_51266 |
| *Enterococcus* | *faecalis* | Enterococcus_faecalis_ATCC_4200 |
| *Enterococcus* | *faecium* | Enterococcus_faecium_ATCC_8459 |
| *Enterococcus* | *saccharolyticus* | Enterococcus_saccharolyticus_ATCC_43076 |
| *Erysipelotrichaceae* | *bacterium* | Erysipelotrichaceae_bacterium_2_2_44A |
| *Escherichia* | *coli* | Escherichia_coli_ATCC_8739 |
| *Escherichia* | *coli* | Escherichia_coli_BW25113 |
| *Escherichia* | *coli* | Escherichia_coli_NC101 |
| *Escherichia* | *coli* | Escherichia_coli_O5_K4_L_H4_str_ATCC_23502 |
| *Escherichia* | *coli* | Escherichia_coli_str_K_12_substr_MG1655 |
| *Escherichia* | *fergusonii* | Escherichia_fergusonii_ATCC_35469 |
| *Eubacterium* | *limosum* | Eubacterium_limosum_DSM_20543 |
| *Eubacterium* | *rectale* | Eubacterium_rectale_ATCC_33656 |
| *Eubacterium* | *ventriosum* | Eubacterium_ventriosum_ATCC_27560 |
| *Faecalibacterium* | *prausnitzii* | Faecalibacterium_prausnitzii_A2_165 |
| *Flavonifractor* | *plautii* | Flavonifractor_plautii_ATCC_29863 |
| *Fusobacterium* | *gonidiaformans* | Fusobacterium_gonidiaformans_ATCC_25563 |
| *Fusobacterium* | *nucleatum* | Fusobacterium_nucleatum_subsp_animalis_ATCC_51191 |
| *Fusobacterium* | *ulcerans* | Fusobacterium_ulcerans_ATCC_49185 |
| *Gardnerella* | *vaginalis* | Gardnerella_vaginalis_JCP7672 |
| *Gemella* | *haemolysans* | Gemella_haemolysans_ATCC_10379 |
| *Gemmiger* | *formicilis* | Gemmiger_formicilis_ATCC_27749 |
| *Granulicatella* | *adiacens* | Granulicatella_adiacens_ATCC_49175 |
| *Haemophilus* | *aegyptius* | *Haemophilus_aegyptius*_ATCC_11116 |
| *Haemophilus* | *haemolyticus* | *Haemophilus_haemolyticus*_HK386 |
| *Helicobacter* | *bilis* | *Helicobacter_bilis*_ATCC_43879 |
| *Helicobacter* | *pylori* | *Helicobacter_pylori*_26695 |
| *Helicobacter* | *rodentium* | *Helicobacter_rodentium*_ATCC_700285 |
| *Holdemania* | *filiformis* | *Holdemania_filiformis*_VPI_J1_31B_1_DSM_12042 |
| *Intestinimonas* | *butyriciproducens* | *Intestinimonas_butyriciproducens*_AF211 |
| *Intestinimonas* | *massiliensis* | *Intestinimonas_massiliensis*_DSM_100417 |
| *Klebsiella* | *michiganensis* | *Klebsiella_michiganensis*_M1 |
| *Klebsiella* | *pneumoniae* | *Klebsiella_pneumoniae*_ATCC_25955 |
| *Klebsiella* | *variicola* | *Klebsiella_variicola*_KV321_ZB100000 |
| *Lachnospiraceae* | *bacterium* | *Lachnospiraceae_bacterium*_1_1_57FAA |
| *Lactobacillus* | *amylolyticus* | *Lactobacillus_amylolyticus*_DSM_11664 |
| *Lactobacillus* | *brevis* | *Lactobacillus_brevis*_ATCC_367 |
| *Lactobacillus* | *casei* | *Lactobacillus_casei*_ATCC_334 |
| *Lactobacillus* | *curvatus* | *Lactobacillus_curvatus*_JCM_1096_DSM_20019 |
| *Lactobacillus* | *fermentum* | *Lactobacillus_fermentum*_ATCC_14931 |
| *Lactobacillus* | *gasseri* | *Lactobacillus_gasseri*_ATCC_33323 |
| *Lactobacillus* | *iners* | *Lactobacillus_iners*_ATCC_55195 |
| *Lactobacillus* | *plantarum* | *Lactobacillus_plantarum_subsp_plantarum*_ATCC_14917 |
| *Lactobacillus* | *reuteri* | *Lactobacillus_reuteri*_ATCC_53608 |
| *Lactobacillus* | *rhamnosus* | *Lactobacillus_rhamnosus*_ATCC_8530 |
| *Lactobacillus* | *ruminis* | *Lactobacillus_ruminis*_ATCC_25644 |
| *Lactobacillus* | *sakei* | *Lactobacillus_sakei_subsp_sakei*_DSM_20017_JCM_1157 |
| *Lactococcus* | *lactis* | *Lactococcus_lactis_subsp_cremoris*_MG1363 |
| *Listeria* | *monocytogenes* | *Listeria_monocytogenes*_ATCC_19117 |
| *Longicatena* | *caecimuris* | *Longicatena_caecimuris*_DSM_29481 |
| *Marvinbryantia* | *formatexigens* | *Marvinbryantia_formatexigens*_I_52_DSM_14469 |
| *Massilibacterium* | *senegalense* | *Massilibacterium_senegalense*_DSM_100455 |
| *Massilioclostridium* | *coli* | *Massilioclostridium_coli*_DSM_103344 |
| *Mediterraneibacter* | *massiliensis* | *Mediterraneibacter_massiliensis*_DSM_100837 |
| *Megasphaera* | *elsdenii* | *Megasphaera_elsdenii*_DSM_20460 |
| *Methanobrevibacter* | *smithii* | *Methanobrevibacter_smithii*_ATCC_35061 |
| *Methanosphaera* | *stadtmanae* | *Methanosphaera_stadtmanae*_DSM_3091 |
| *Methylobacterium* | *radiotolerans* | *Methylobacterium_radiotolerans*_JCM_2831 |
| *Micrococcus* | *luteus* | *Micrococcus_luteus*_NCTC_2665 |
| *Mitsuokella* | *multacida* | *Mitsuokella_multacida*_DSM_20544 |
| *Mobilicoccus* | *massiliensis* | *Mobilicoccus_massiliensis*_DSM_29065 |
| *Mobiluncus* | *curtisii* | *Mobiluncus_curtisii*_ATCC_51333 |
| *Mogibacterium* | *timidum* | *Mogibacterium_timidum*_ATCC_33093 |
| *Muricomes* | *intestini* | *Muricomes_intestini*_DSM_29489 |
| *Mycobacterium* | *avium* | *Mycobacterium_avium_subsp_avium*_ATCC_25291 |
| *Mycoplasma* | *pneumoniae* | *Mycoplasma_pneumoniae*_M129_B7 |
| *Neisseria* | *elongata* | *Neisseria_elongata_subsp_glycolytica*_ATCC_29315 |
| *Neisseria* | *mucosa* | *Neisseria_mucosa*_ATCC_25996 |
| *Neofamilia* | *massiliensis* | *Neofamilia_massiliensis*_DSM_100639 |
| *Nevskia* | *ramosa* | *Nevskia_ramosa*_DSM_11499 |
| *Novosphingobium* | *aromaticivorans* | *Novosphingobium_aromaticivorans*_DSM_12444 |
| *Obesumbacterium* | *proteus* | *Obesumbacterium_proteus*_ATCC_12841 |
| *Olegusella* | *massiliensis* | *Olegusella_massiliensis*_DSM_101849 |
| *Olsenella* | *uli* | *Olsenella_uli*_DSM_7084 |
| *Paenibacillus* | *alvei* | *Paenibacillus_alvei*_DSM_29 |
| *Paenibacillus* | *ihumii* | *Paenibacillus_ihumii*_DSM_100664 |
| *Paenibacillus* | *tuaregi* | *Paenibacillus_tuaregi*_DSM_102801 |
| *Parabacteroides* | *chinchillae* | *Parabacteroides_chinchillae*_DSM_29073 |
| *Parabacteroides* | *distasonis* | *Parabacteroides_distasonis*_ATCC_8503 |
| *Parabacteroides* | *goldsteinii* | *Parabacteroides_goldsteinii*_DSM_19448_WAL_12034 |
| *Parabacteroides* | *gordonii* | *Parabacteroides_gordonii*_DSM_23371 |
| *Parabacteroides* | *johnsonii* | *Parabacteroides_johnsonii*_DSM_18315 |
| *Parabacteroides* | *merdae* | *Parabacteroides_merdae*_ATCC_43184 |
| *Paraclostridium* | *bifermentans* | *Paraclostridium_bifermentans*_ATCC_638 |
| *Pasteurella* | *caecimuris* | *Pasteurella_caecimuris*_NM44_TS2_9 |
| *Pediococcus* | *acidilactici* | *Pediococcus_acidilactici*_DSM_20284 |
| *Peptoniphilus* | *duerdenii* | *Peptoniphilus_duerdenii*_ATCC_BAA_1640 |
| *Peptostreptococcus* | *anaerobius* | *Peptostreptococcus_anaerobius*_DSM_2949 |
| *Phascolarctobacterium* | *faecium* | *Phascolarctobacterium_faecium*_DSM_14760 |
| *Porphyromonas* | *gingivalis* | *Porphyromonas_gingivalis*_ATCC_33277 |
| *Prevotella* | *albensis* | *Prevotella_albensis_DSM_11370* |
| *Prevotella* | *buccae* | *Prevotella_buccae_ATCC_33574* |
| *Prevotella* | *buccalis* | *Prevotella_buccalis_ATCC_35310* |
| *Prevotella* | *copri* | *Prevotella_copri_CB7_DSM_18205* |
| *Prevotella* | *melaninogenica* | *Prevotella_melaninogenica_ATCC_25845* |
| *Propionibacterium* | *acidifaciens* | *Propionibacterium_acidifaciens_DSM_21887* |
| *Propionibacterium* | *jensenii* | *Propionibacterium_jensenii_DSM_20535* |
| *Proteus* | *mirabilis* | *Proteus_mirabilis_ATCC_29906* |
| *Proteus* | *vulgaris* | *Proteus_vulgaris_ATCC_49132* |
| *Providencia* | *rettgeri* | *Providencia_rettgeri_DSM_1131* |
| *Providencia* | *stuartii* | *Providencia_stuartii_ATCC_25827* |
| *Pseudomonas* | *aeruginosa* | *Pseudomonas_aeruginosa_PAO1* |
| *Pseudomonas* | *stutzeri* | *Pseudomonas_stutzeri_DSM_4166* |
| *Raoultella* | *planticola* | *Raoultella_planticola_ATCC_33531* |
| *Rhodococcus* | *rhodochrous* | *Rhodococcus_rhodochrous_ATCC_17895* |
| *Rikenella* | *microfus* | *Rikenella_microfus_DSM_15922* |
| *Roseburia* | *inulinivorans* | *Roseburia_inulinivorans_DSM_16841* |
| *Roseomonas* | *mucosa* | *Roseomonas_mucosa_ATCC_BAA_692* |
| *Ruminiclostridium* | *thermocellum* | *Ruminiclostridium_thermocellum_ATCC_27405* |
| *Ruminococcus* | *bromii* | *Ruminococcus_bromii_ATCC_27255* |
| *Ruminococcus* | *flavifaciens* | *Ruminococcus_flavifaciens_ATCC_19208* |
| *Ruminococcus* | *gauvreauii* | *Ruminococcus_gauvreauii_DSM_19829* |
| *Ruminococcus* | *gnavus* | *Ruminococcus_gnavus_ATCC_29149* |
| *Ruminococcus* | *torques* | *Ruminococcus_torques_ATCC_27756* |
| *Salmonella* | *bongori* | *Salmonella_bongori_NCTC_12419* |
| *Salmonella* | *enterica* | *Salmonella_enterica_subsp_enterica_serovar_Typhimurium_ATCC_14028* |
| *Selenomonas* | *bovis* | *Selenomonas_bovis_DSM_23594* |
| *Selenomonas* | *sputigena* | *Selenomonas_sputigena_ATCC_35185* |
| *Serratia* | *marcescens* | *Serratia_marcescens_LCT_SM213* |
| *Shewanella* | *woodyi* | *Shewanella_woodyi_ATCC_51908* |
| *Shigella* | *flexneri* | *Shigella_flexneri_CDC_796_83* |
| *Shuttleworthia* | *satelles* | *Shuttleworthia_satelles_DSM_14600* |
| *Silanimonas* | *lenta* | *Silanimonas_lenta_DSM_16282* |
| *Sinobacterium* | *caligoides* | *Sinobacterium_caligoides_DSM_100316* |
| *Slackia* | *equolifaciens* | *Slackia_equolifaciens_DSM_24851* |
| *Slackia* | *exigua* | *Slackia_exigua_ATCC_700122* |
| *Solobacterium* | *moorei* | *Solobacterium_moorei_DSM_22971* |
| *Sphingosinicella* | *microcystinivorans* | *Sphingosinicella_microcystinivorans_DSM_19791* |
| *Staphylococcus* | *aureus* | *Staphylococcus_aureus_subsp_aureus_ATCC_51811* |
| *Staphylococcus* | *haemolyticus* | *Staphylococcus_haemolyticus_JCSC1435* |
| *Staphylococcus* | *pasteuri* | *Staphylococcus_pasteuri_SP1* |
| *Staphylococcus* | *saprophyticus* | *Staphylococcus_saprophyticus_subsp_saprophyticus_ATCC_15305* |
| *Stenotrophomonas* | *maltoiphilia* | *Stenotrophomonas_maltoiphilia_ATCC_19867* |
| *Stenotrophomonas* | *rhizophila* | *Stenotrophomonas_rhizophila_PCA13* |
| *Streptococcus* | *australis* | *Streptococcus_australis_ATCC_700641* |
| *Streptococcus* | *bovis* | *Streptococcus_bovis_ATCC_700338* |
| *Streptococcus* | *cristatus* | *Streptococcus_cristatus_ATCC_51100* |
| *Streptococcus* | *infantis* | *Streptococcus_infantis_ATCC_700779* |
| *Streptococcus* | *mutans* | *Streptococcus_mutans_ATCC_25175* |
| *Streptococcus* | *oralis* | *Streptococcus_oralis_ATCC_35037* |
| *Streptococcus* | *parasanguinis* | *Streptococcus_parasanguinis_ATCC_903* |
| *Streptococcus* | *peroris* | *Streptococcus_peroris_ATCC_700780* |
| *Streptococcus* | *pyogenes* | *Streptococcus_pyogenes_ATCC_10782* |
| *Streptococcus* | *sanguinis* | *Streptococcus_sanguinis_ATCC_29667* |
| *Subdoligranulum* | *variabile* | *Subdoligranulum_variabile_DSM_15176* |
| *Tatumella* | *ptyseos* | *Tatumella_ptyseos_ATCC_33301* |
| *Timonella* | *senegalensis* | *Timonella_senegalensis_JC301* |
| *Trabulsiella* | *guamensis* | *Trabulsiella_guamensis_ATCC_49490* |
| *Treponema* | *denticola* | *Treponema_denticola_ATCC_33520* |
| *Treponema* | *socranskii* | *Treponema_socranskii_subsp_socranskii_VPI_DR56BR1116_ATCC_35536* |
| *Ureaplasma* | *parvum* | *Ureaplasma_parvum_serovar_1_str_ATCC_27813* |
| *Vagococcus* | *fluvialis* | *Vagococcus_fluvialis_DSM_5731* |
| *Varibaculum* | *cambriense* | *Varibaculum_cambriense_DSM_15806* |
| *Variovorax* | *paradoxus* | *Variovorax_paradoxus_B4* |
| *Veillonella* | *denticariosi* | *Veillonella_denticariosi_JCM_15641* |
| *Veillonella* | *dispar* | *Veillonella_dispar_ATCC_17748* |
| *Veillonella* | *parvula* | *Veillonella_parvula_ATCC_17745* |
| *Vibrio* | *campbellii* | Vibrio_campbellii_ATCC_BAA_1116 |
| *Vibrio* | *cholerae* | Vibrio_cholerae_A1552 |
| *Vibrio* | *sinaloensis* | Vibrio_sinaloensis_DSM_21326 |
| *Weissella* | *paramesenteroides* | Weissella_paramesenteroides_ATCC_33313 |
| *Yersinia* | *bercoviei* | Yersinia_bercoviei_ATCC_43970 |
| *Yersinia* | *kristensenii* | Yersinia_kristensenii_ATCC_33638 |

## REFERENCES

1. Jun, S., Si, F., Pugatch, R. & Scott, M. Fundamental principles in bacterial physiology—history, recent progress, and the future with focus on cell size control: a review. Rep. Prog. Phys. 81, 056601 (2018).

2. Xu, L., Zakem, E. & Weissman, J. L. Improved maximum growth rate prediction from microbial genomes by integrating phylogenetic information. Nat Commun 16, 4256 (2025).

3. Ramon, C. & Stelling, J. Functional comparison of metabolic networks across species. Nat Commun 14, 1699 (2023).

4. Kanehisa, M. & Goto, S. KEGG: Kyoto Encyclopedia of Genes and Genomes. Nucleic Acids Res 28, 27–30 (2000).

5. Nielsen, J. & Keasling, J. D. Engineering cellular metabolism. Cell 164, 1185–1197 (2016).

6. Ke, J., Wang, B. & Yoshikuni, Y. Microbiome engineering: synthetic biology of plant-associated microbiomes in sustainable agriculture. Trends in Biotechnology 39, 244–261 (2021).

7. Monod, J. The growth of bacterial cultures. Annu Rev Microbiol 3, 371–394 (1949).

8. Schaechter, M., MaalØe, O. & Kjeldgaard, N. O. Dependency on medium and temperature of cell size and chemical composition during balanced growth of *Salmonella typhimurium*. Microbiology 19, 592–606 (1958).

9. Neidhardt, F. C. Escherichia Coli and Salmonella Typhimurium : Cellular and Molecular Biology. (American Society for Microbiology, Washington, D.C, 1987).

10. Chubukov, V., Gerosa, L., Kochanowski, K. & Sauer, U. Coordination of microbial metabolism. Nat Rev Microbiol 12, 327–340 (2014).

11. Scott, M. & Hwa, T. Bacterial growth laws and their applications. Current Opinion in Biotechnology 22, 559–565 (2011).

12. Orth, J. D., Thiele, I. & Palsson, B. Ø. What is flux balance analysis? Nat Biotechnol 28, 245–248 (2010).

13. Gu, C., Kim, G. B., Kim, W. J., Kim, H. U. & Lee, S. Y. Current status and applications of genome-scale metabolic models. Genome Biol 20, 121 (2019).

14. Okano, H., Hermsen, R., Kochanowski, K. & Hwa, T. Regulation underlying hierarchical and simultaneous utilization of carbon substrates by flux sensors in Escherichia coli. Nat Microbiol 5, 206–215 (2020).

15. Hermsen, R., Okano, H., You, C., Werner, N. & Hwa, T. A growth-rate composition formula for the growth of *E. coli* on co-utilized carbon substrates. Mol Syst Biol 11, 801 (2015).

16. Scott, M., Gunderson, C. W., Mateescu, E. M., Zhang, Z. & Hwa, T. Interdependence of cell growth and gene expression: origins and consequences. Science 330, 1099–1102 (2010).

17. Hui, S. et al. Quantitative proteomic analysis reveals a simple strategy of global resource allocation in bacteria. Mol Syst Biol 11, MSB145697 (2015).

18. Mehta, P. A twenty-first century statistical physics of life. arXiv, preprint at 10.48550/arXiv.2410.20506 (2024).

19. Youk, H. & van Oudenaarden, A. Growth landscape formed by perception and import of glucose in yeast. Nature 462, 875–879 (2009).

20. Litsios, A., Ortega, Á. D., Wit, E. C. & Heinemann, M. Metabolic-flux dependent regulation of microbial physiology. Curr Opin Microbiol 42, 71–78 (2018).

21. Jeong, H., Tombor, B., Albert, R., Oltvai, Z. N. & Barabási, A.-L. The large-scale organization of metabolic networks. Nature 407, 651–654 (2000).

22. Maslov, S., Krishna, S., Pang, T. Y. & Sneppen, K. Toolbox model of evolution of prokaryotic metabolic networks and their regulation. Proc Natl Acad Sci 106, 9743–9748 (2009).

23. Noor, E., Eden, E., Milo, R. & Alon, U. Central carbon metabolism as a minimal biochemical walk between precursors for biomass and energy. Mol Cell 39, 809–820 (2010).

24. Scott, M., Klumpp, S., Mateescu, E. M. & Hwa, T. Emergence of robust growth laws from optimal regulation of ribosome synthesis. Mol Syst Biol 10, MSB145379 (2014).

25. Bren, A. et al. Glucose becomes one of the worst carbon sources for E.coli on poor nitrogen sources due to suboptimal levels of cAMP. Sci Rep 6, 24834 (2016).

26. Perrin, E. et al. Diauxie and co-utilization of carbon sources can coexist during bacterial growth in nutritionally complex environments. Nat Commun 11, 3135 (2020).

27. Bakkeren, E., Piskovsky, V. & Foster, K. R. Metabolic ecology of microbiomes: Nutrient competition, host benefits, and community engineering. Cell Host & Microbe 33, 790–807 (2025).

28. Zhu, M., Mori, M., Hwa, T. & Dai, X. Distantly related bacteria share a rigid proteome allocation strategy with flexible enzyme kinetics. Proc Natl Acad Sci 122, e2427091122 (2025).

29. Ashburner, M. et al. Gene Ontology: tool for the unification of biology. Nat Genet 25, 25–29 (2000).

30. Salgado, H. et al. RegulonDB v12.0: a comprehensive resource of transcriptional regulation in E. coli K-12. Nucleic Acids Res 52, D255–D264 (2024).

31. Sastry, A. V. et al. The *Escherichia coli* transcriptome mostly consists of independently regulated modules. Nat Commun 10, 5536 (2019).

32. Schuster, S., Fell, D. A. & Dandekar, T. A general definition of metabolic pathways useful for systematic organization and analysis of complex metabolic networks. Nat Biotechnol 18, 326–332 (2000).

33. Chan, S. H. J. & Ji, P. Decomposing flux distributions into elementary flux modes in genome-scale metabolic networks. Bioinformatics 27, 2256–2262 (2011).

34. Mori, M., Cheng, C., Taylor, B. R., Okano, H. & Hwa, T. Functional decomposition of metabolism allows a system-level quantification of fluxes and protein allocation towards specific metabolic functions. Nat Commun 14, 4161 (2023).

35. Ho, P.-Y., Nguyen, T. H., Sanchez, J. M., DeFelice, B. C. & Huang, K. C. Resource competition predicts assembly of gut bacterial communities in vitro. Nat Microbiol 9, 1036–1048 (2024).

36. Towbin, B. D. et al. Optimality and sub-optimality in a bacterial growth law. Nat Commun 8, 14123 (2017).

37. Heinken, A. et al. Genome-scale metabolic reconstruction of 7,302 human microorganisms for personalized medicine. Nat Biotechnol 41, 1320–1331 (2023).

38. Baldazzi, V., Ropers, D., Gouzé, J.-L., Gedeon, T. & de Jong, H. Resource allocation accounts for the large variability of rate-yield phenotypes across bacterial strains. eLife 12, e79815 (2023).

39. 39. Xiong, Z. J., Huisman, J. S. & Gore, J. Growth rate and yield are not constrained by a trade-off in microbes. bioRxiv, Preprint at 10.64898/2026.09.08.750265 (2026).

40. Price, N. D., Reed, J. L. & Palsson, B. Ø. Genome-scale models of microbial cells: evaluating the consequences of constraints. Nat Rev Microbiol 2, 886–897 (2004).

41. Schuetz, R., Kuepfer, L. & Sauer, U. Systematic evaluation of objective functions for predicting intracellular fluxes in *Escherichia coli*. Mol Syst Biol 3, MSB4100162 (2007).

42. O’Brien, E. J., Lerman, J. A., Chang, R. L., Hyduke, D. R. & Palsson, B. Ø. Genome-scale models of metabolism and gene expression extend and refine growth phenotype prediction. Mol Syst Biol 9, MSB201352 (2013).

43. Klamt, S., Müller, S., Regensburger, G. & Zanghellini, J. A mathematical framework for yield (*vs.* rate) optimization in constraint-based modeling and applications in metabolic engineering. Metab Eng 47, 153–169 (2018).

44. Heinken, A. et al. A genome-scale metabolic reconstruction resource of 247,092 diverse human microbes spanning multiple continents, age groups, and body sites. Cell Syst 16, 101196 (2025).

45. King, Z. A. et al. BiGG Models: A platform for integrating, standardizing and sharing genome-scale models. Nucleic Acids Res 44, D515–D522 (2016).

46. Basan, M. et al. Overflow metabolism in *Escherichia coli* results from efficient proteome allocation. Nature 528, 99–104 (2015).

47. Takano, S., Vila, J. C. C., Miyazaki, R., Sánchez, Á. & Bajić, D. The architecture of metabolic networks constrains the evolution of microbial resource hierarchies. Mol Biol Evol 40, msad187 (2023).

48. Fritzemeier, C. J., Hartleb, D., Szappanos, B., Papp, B. & Lercher, M. J. Erroneous energy-generating cycles in published genome scale metabolic networks: Identification and removal. PLOS Comput Biol 13, e1005494 (2017).

49. Beguerisse-Díaz, M., Bosque, G., Oyarzún, D., Picó, J. & Barahona, M. Flux-dependent graphs for metabolic networks. Npj Syst Biol Appl 4, 1–14 (2018).

50. Grover, A. & Leskovec, J. node2vec: scalable feature learning for networks. in Proceedings of the 22nd ACM SIGKDD International Conference on Knowledge Discovery and Data Mining 855–864 (ACM, San Francisco California USA, 2016). doi:10.1145/2939672.2939754.

51. Zamboni, N., Mouncey, N., Hohmann, H.-P. & Sauer, U. Reducing maintenance metabolism by metabolic engineering of respiration improves riboflavin production by *Bacillus subtilis*. Metab Eng 5, 49–55 (2003).

52. Ricaurte, D. et al. High-throughput transcriptomics of 409 bacteria–drug pairs reveals drivers of gut microbiota perturbation. Nat Microbiol 9, 561–575 (2024).

53. Bekiaris, P. S. & Klamt, S. Automatic construction of metabolic models with enzyme constraints. BMC Bioinform 21, 19 (2020).

54. Beg, Q. K. et al. Intracellular crowding defines the mode and sequence of substrate uptake by *Escherichia coli* and constrains its metabolic activity. Proc Natl Acad Sci 104, 12663–12668 (2007).

55. Yamagishi, J. F. & Hatakeyama, T. S. Global constraint principle for microbial growth laws. Proc Natl Acad Sci 122, e2515031122 (2025).

56. Mori, M. et al. From coarse to fine: the absolute *Escherichia coli* proteome under diverse growth conditions. Mol Syst Biol 17, MSB20209536 (2021).

57. Balakrishnan, R. et al. Principles of gene regulation quantitatively connect DNA to RNA and proteins in bacteria. Science 378, eabk2066 (2022).

58. Ramseier, T. M. Cra and the control of carbon flux via metabolic pathways. Res Microbiol 147, 489–493 (1996).

59. Cooper, R. A. & Kornberg, H. L. Net formation of phosphoenolpyruvate from pyruvate by *Escherichia coli*. Biochimica et Biophysica Acta (BBA) - General Subjects 104, 618–620 (1965).

60. Dudek, C.-A. & Jahn, D. PRODORIC: state-of-the-art database of prokaryotic gene regulation. Nucleic Acids Res 50, D295–D302 (2022).

61. Caldara, M. et al. Arginine biosynthesis in *Escherichia coli*. J Biol Chem 283, 6347–6358 (2008).

62. Lu, C.-D., Yang, Z. & Li, W. Transcriptome analysis of the ArgR regulon in *Pseudomonas aeruginosa*. J Bacteriol 186, 3855–3861 (2004).

63. Wu, C. et al. Cellular perception of growth rate and the mechanistic origin of bacterial growth law. Proc Natl Acad Sci 119, e2201585119 (2022).

64. Thiermann, R. et al. Decoupling of global metabolic flux and proteome partitioning in bacteria. Science 392, eaeb6410 (2026).

65. Mueller, F. M. et al. Non-canonical resource allocation in heterotrophically growing *Thermoanaerobacter kivui*. Nat Commun 16, 8489 (2025).

66. Massucci, F. A., Guimerà, R., Nunes Amaral, L. A. & Sales-Pardo, M. Scaling and optimal synergy: Two principles determining microbial growth in complex media. Phys Rev E 91, 062703 (2015).

67. 67. Tilman, D. Resource Competition and Community Structure*. (MPB-17), Volume 17*. (Princeton University Press, 1982).

68. Huelsmann, M., Schubert, O. T. & Ackermann, M. A framework for understanding collective microbiome metabolism. Nat Microbiol 9, 3097–3109 (2024).

69. Li, L. et al. Revealing proteome-level functional redundancy in the human gut microbiome using ultra-deep metaproteomics. Nat Commun 14, 3428 (2023).

70. Alzoubi, D., Desouki, A. A. & Lercher, M. J. Flux balance analysis with or without molecular crowding fails to predict two thirds of experimentally observed epistasis in yeast. Sci Rep 9, 11837 (2019).

71. Chen, Y. et al. Reconstruction, simulation and analysis of enzyme-constrained metabolic models using GECKO Toolbox 3.0. Nat Protoc 19, 629–667 (2024).

72. Adadi, R., Volkmer, B., Milo, R., Heinemann, M. & Shlomi, T. Prediction of microbial growth rate versus biomass yield by a metabolic network with kinetic parameters. PLOS Comput Biol 8, e1002575 (2012).

73. Lewis, N. E. et al. Omic data from evolved *E. coli* are consistent with computed optimal growth from genome-scale models. Mol Syst Biol 6, MSB201047 (2010).

74. Ebrahim, A., Lerman, J. A., Palsson, B. O. & Hyduke, D. R. COBRApy: COnstraints-Based Reconstruction and Analysis for Python. BMC Syst Biol 7, 74 (2013).

75. Gurobi Optimization, LLC. Gurobi Optimizer Reference Manual. (2026).

76. MacQueen, J. Some methods for classification and analysis of multivariate observations. in Proceedings of the Fifth Berkeley Symposium on Mathematical Statistics and Probability, Volume 1: Statistics vol. 5.1 281–298 (University of California Press, 1967).

77. Henry, C. S. et al. High-throughput generation, optimization and analysis of genome-scale metabolic models. Nat Biotechnol 28, 977–982 (2010).

78. Aziz, R. K. et al. The RAST Server: Rapid Annotations using Subsystems Technology. BMC Genom 9, 75 (2008).

79. Benjamini, Y. & Hochberg, Y. Controlling the false discovery rate: a practical and powerful approach to multiple testing. Royal Statistical Society. Journal. Series B: Methodological 57, 289–300 (1995).

80. Darzi, Y., Letunic, I., Bork, P. & Yamada, T. iPath3.0: interactive pathways explorer v3. Nucleic Acids Res 46, W510–W513 (2018).

